# Multi-scale phenological niches of seed fall in diverse Amazonian plant communities

**DOI:** 10.1101/2021.04.10.438715

**Authors:** Damie Pak, Varun Swamy, Patricia Alvarez-Loayza, Fernando Cornejo, Simon A. Queenborough, Margaret R. Metz, John Terborgh, Renato Valencia, S. Joseph Wright, Nancy C. Garwood, Jesse R. Lasky

## Abstract

Phenology has long been hypothesized as an avenue for niche partitioning or interspecific facilitation, both promoting species coexistence. Tropical plant communities exhibit striking diversity in reproductive phenology, but many are also noted for large synchronous reproductive events. Here we study whether the phenology of seed fall in such communities is non-random, what are the temporal scales of phenological patterns, and ecological factors that drive reproductive phenology. We applied multivariate wavelet analyses to test for phenological synchrony versus compensatory dynamics (*i.e*. anti-synchronous patterns where one species’ decline is compensated by the rise of another) among species and across temporal scales. We used data from long-term seed rain monitoring of hyperdiverse plant communities in the western Amazon. We found significant synchronous whole-community phenology at multiple time scales, consistent with shared environmental responses or positive interactions among species. We also observed both compensatory and synchronous phenology within groups of species likely to share traits (confamilials) and seed dispersal mechanisms. Wind-dispersed species exhibited significant synchrony at ~6 mo scales, suggesting these species share phenological niches to match seasonality of wind. Our results suggest that community phenology is shaped by shared environmental responses but that the diversity of tropical plant phenology may partly result from temporal niche partitioning. The scale-specificity and time-localized nature of community phenology patterns highlights the importance of multiple and shifting drivers of phenology.

**Open research statement:** Data are provided as private-for-peer review. Code and data can be found at https://github.com/pakdamie/treephenology which is currently public.

This submission does not use novel code.

## INTRODUCTION

Species within ecological communities often exhibit interspecific diversity in the phenology of key life events. This diversity may represent an axis of niche partitioning that reflects community assembly and evolutionary history (Ashton et al. 1988, Gonzalez and Loreau 2009, Wolkovich and Cleland 2011, Bernard-Verdier et al. 2012, Godoy and Levine 2013). Species differences in phenology may limit interspecific competition and promote species coexistence by causing niche complementarity through time in resource use or interactions with mutualists like pollinators, or in apparent competition mediated by natural enemies (Robertson 1895, Rathcke and Lacey 1985). Alternatively, phenological synchrony may be promoted by pulses in resource supply or periodically harsh environmental conditions that limit the possible phenological options (Gentry 1974, Rathcke and Lacey 1985, Vasseur et al. 2014, Usinowicz et al. 2017, Detto et al. 2018). Facilitation due to enhanced attraction of mutualist animals or predator satiation may also promote synchronous reproduction (Janzen 1974). However, phenology remains a relatively poorly characterized dimension of functional diversity in many communities, owing to a lack of long-term monitoring and the multi-scale complexity of phenology (Wolkovich et al. 2014).

Co-occurring plants with similar reproductive phenology might be more likely to compete for frugivores (Saracco et al. 2005) or other resources, given the requirements of reproduction (Karlsson and Méndez 2005). As a result, those species capable of coexisting might partition phenological space. Researchers have studied phenological niches in tropical forests (e.g. Gentry 1974, Stiles 1977, Wheelwright 1985, Ashton et al. 1988, Poulin et al. 1999, Jones and Comita 2010) and elsewhere (Elzinga et al. 2007, Botes et al. 2008, Albrecht et al. 2015). Within a community, phenological niche partitioning might be strongest among species with shared mutualists or shared resource requirements (Encinas-Viso et al. 2012), similarities that often characterize phylogenetically related species (Robertson 1895, Prinzing et al. 2001, Donoghue 2008, Davies et al. 2013). However, there are simultaneous and opposing processes interacting with phenology, such as abiotic seasonality versus resource competition. As a result, phenological patterns indicative of shared temporal niches (interspecific synchrony) versus temporal niche partitioning (interspecific compensation, or anti-synchrony) may only emerge at certain time scales or over certain periods of time (Baird 1980, Vasseur et al. 2005, Keitt 2008, Lasky et al. 2016).

Tropical plant communities have highly varied phenology and there are often multiple species reproducing at any given time of the year (Frankie et al. 1974, Gentry 1974, van Schaik et al. 1993), including for the stage that we study here: seed fall (Smythe 1970, Chang-Yang et al. 2016, Detto et al. 2018). The phenological diversity of tropical plants may be made possible by favorable temperature and (in rainforests) moisture for much of the year (Gentry 1974, Usinowicz et al. 2017). Despite this potential, tropical plant communities often do exhibit synchrony among at least a subset of the community, perhaps due to shared responses to abiotic seasonality (van Schaik et al. 1993, Detto et al. 2018) or seasonality in frugivory and seed dispersal (Poulin et al. 1999). It seems likely that different species may be limited by different conditions fluctuating across the year (e.g. light, moisture, heat), thus diversity in the phenology of tropical seed fall may be a consequence of distinct strategies or sensitivities to seasonality in resources (Lasky et al. 2016). Additionally, positive density dependent interactions among species may promote synchrony, for example masting that overwhelms seed predators (Ashton et al. 1988, Jones and Comita 2010) or facilitation by frugivore attraction (Carlo 2005). Abiotic conditions may control species interactions, such that synchronous or compensatory reproduction occur during periods with specific abiotic conditions (Vasseur et al. 2005).

We used wavelet analyses to determine whether species exhibited synchronous or compensatory (anti-synchronous) seed rain (illustrated in Figure 1) (Lasky et al. 2016). Wavelets are basis functions, combinations of which can be used to characterize signals in data (here, time series of seed rain). Wavelet transformations decompose signals into patterns at different scales, like other spectral analyses, but with the added advantage that wavelets can characterize time-localized and nonstationary patterns, i.e. patterns that are inconsistent over a time series (Terrence and Compo 1998, Keitt 2008). In the wavelet transformation, the base wavelet is translated across the time series at varying scales/frequencies of the wavelet to identify the important time scales that contribute to the variability in the signal (Cazelles et al. 2008). By resolving non-stationary and scale-specific patterns, we may improve our ability to detect multiple opposing processes affecting seed rain dynamics at different temporal scales (scale-specific) or points in time (non-stationary). For example, while species may all increase reproduction during once-a-year seasons of high resource supply (annual-scale synchrony), species may peak in reproduction at different points within a favorable season (Figure 1, within-season-scale compensatory dynamics, Lasky et al. 2016).

**Figure 1.**
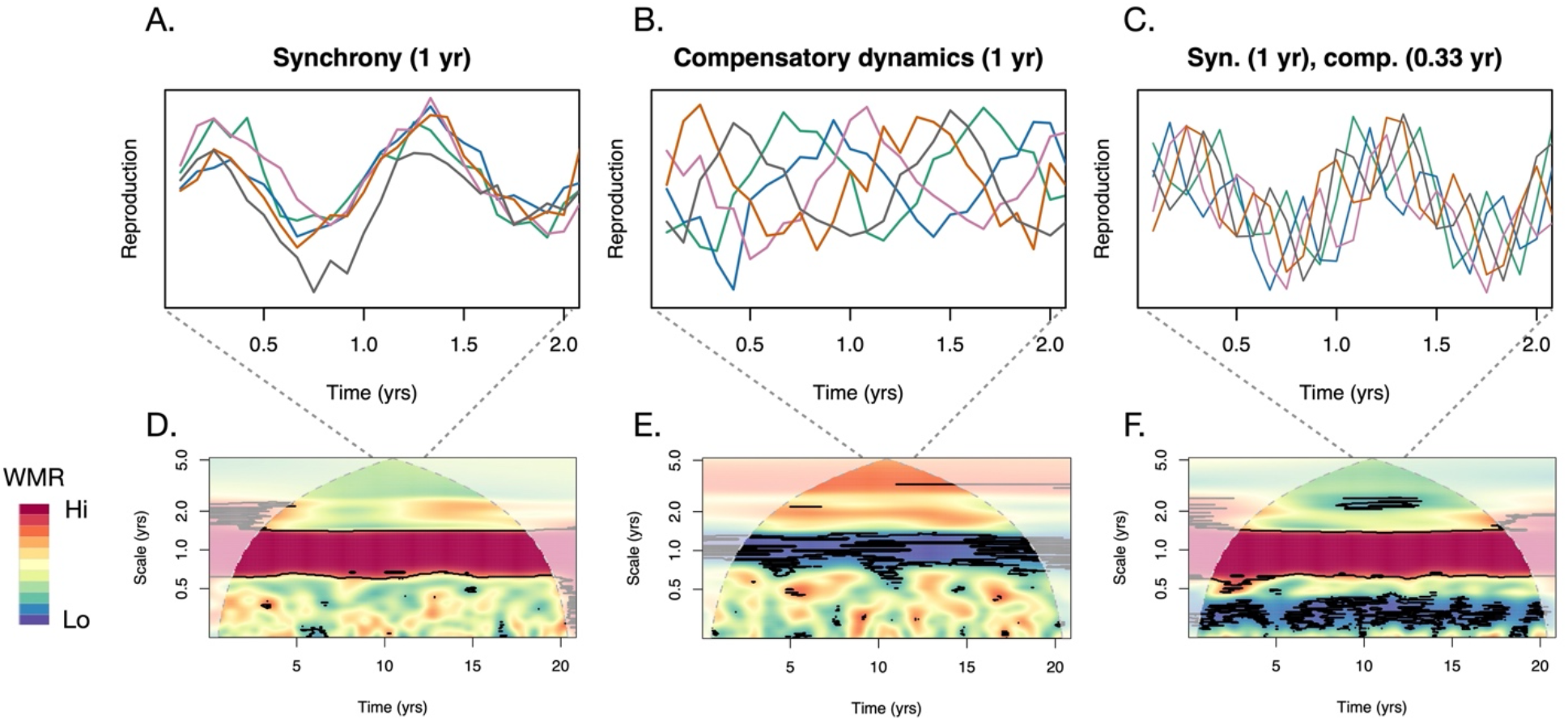
Illustration of wavelet modulus ratio (WMR) as a measure of simulated multi-species patterns in reproduction. (A-C) Shows two-year time series of reproduction (excerpted from a 20-year series) for three different scenarios for a 5-species community (each species a different colored line). (D-F) Demonstrates WMR for these scenarios across a whole 20-year time series, where WMR is determined by the variation in total aggregate community reproduction relative to the variation in individual species reproduction. Thick black lines indicate regions of significantly non-random WMR (FDR = 0.05). When there is synchrony (A), the aggregate community variation is similar to the individual species variation, and WMR is high at the relevant scale (1 yr). When there are compensatory dynamics (B), the aggregate community variation is low but the individual species variation is high, and WMR is low at the relevant scale (1 yr). Both synchrony and compensatory dynamics can occur at different scales (C), in this case long-scale synchrony and short scale compensation. A small amount of noise was added to each scenario. The cone of influence (white shading in D-F) marks the regions where the wavelet transforms are affected by the boundaries of the sampling period.

We addressed the following questions:

1. Do communities exhibit compensatory patterns or synchronous patterns of reproduction (seed fall) through time?
2. Is evidence for compensatory and synchronous dynamics scale-specific or non-stationary (i.e. changing through time)?
3. Is evidence for niche partitioning of seed fall phenology strongest among functionally similar species, potentially those with the greatest likelihood of interspecific competition? Do related species exhibit stronger compensatory dynamics? Or alternatively do morphologically similar or related species exhibit synchrony?
4. Does community phenology differ between sites and time periods of contrasting climate, suggesting phenological niches may be mediated by fluctuations in environment?

## METHODS

### Study sites

We studied two forest plots in the western Amazon basin, in Cocha Cashu, Peru and Yasuní, Ecuador (Figure S1). These plots were monitored across different intervals, from February 2000–February 2017 in Yasuní and September 2002–January 2011 in Cocha Cashu. The study plot in Ecuador was located in Yasuní National Park at the Estación Científica Yasuní (0° 41’ S, 76° 24’ W), a research station maintained by Pontificia Universidad Católica del Ecuador. The Yasuní lowland rain forest is in the wettest and least seasonal region of the Amazon (Xiao et al. 2006, Silman 2007). Mean annual rainfall is 2826 mm, with no months having <100 mm rainfall on average (Valencia et al. 2004b, 2004a). Seed traps were placed within the 50-ha Yasuní Forest Dynamics Plot (YFDP, established in 1995), where elevation ranged from 216 to 248 m. In 25 ha of this forest plot, 1104 tree species were recorded (Valencia et al. 2004b, 2004a).

The study plot in Peru is located at Cocha Cashu Biological Station (11°54’S, 71°22’W), which is situated at 360 m mean elevation within the core area of Manu National Park, at the western margin of the Madre de Dios river basin. The study plot is located in mature floodplain forest habitat, which comprises over 700 tree species (Pitman et al. 2002). Annual precipitation ranges between 2000–2500 mm, with a pronounced dry season from June to October with typically less than 100 mm monthly rainfall (Gentry 1993). In the period from September and April there is an excess of fruit available for frugivorous vertebrates (Terborgh 1986b), which may indicate that plants compete to attract frugivores during this period.

### Seed rain data

In each plot, an array of seed traps was established. At Yasuní we followed the methods of Wright and Calderon (1995). In February 2000, 200 seed traps were placed in the 50-ha YFDP along trails but >50 m from the plot border. Every 13.5 m along the trails, a trap was placed a random distance between 4 and 10 m perpendicular from the trail, alternating left and right. Traps were constructed of PVC tubes and 1-mm fiberglass mesh, positioned 0.75 m above ground, with an area of 0.57 m^2^. Twice a month (median interval = 20 days, SD = 7.0 days), from February 2000 to February 2017 all reproductive parts in each trap were counted and identified to species or morphospecies using a reference collection of seeds and fruits maintained on site.

At Cocha Cashu, year-round fruit and seed fall were counted between 2002 and 2011 within a 4-ha plot. A 17 × 17 array of 289 evenly spaced seed-fall traps was installed within the central 1.44 ha (120 × 120 m) of the plot at the beginning of the study. Seed traps consisted of 0.49 m^2^ (70 × 70 cm) open bags made of 1-mm nylon mesh sewn to wire frames with 0.5-mm monofilament line. Corners of the traps were attached to nearby trees with 1-mm monofilament line so that the traps were suspended approximately 1 m above the ground. The contents of the traps were collected every ~2 weeks (median interval = 15 days, SD = 3.3 days), and all seeds, fruit and fruit parts (capsules, valves, pods, etc.) were identified to species and recorded.

For fruit counts at both sites we estimated number of seeds collected by multiplying by the average number of seeds per fruit. We divided counts by the number of days in intervals to control for slight variation in intervals. Further detail is available in the Supplemental Material.

### Seed dispersal mechanisms

We grouped species into different dispersal syndromes to test for non-random WMR within each syndrome, using slightly different approaches for each site depending on available data. At each site, we conducted two separate classification efforts, one for all species, and another more granular classification focused on tree species (excluding lianas, herbaceous, and woody shrubs).

At both sites for all plants we first focused our analysis on classifications as animal (Yasuní, N = 741, Cocha Cashu N = 174) or wind (Yasuní N = 139, Cocha Cashu N = 48) dispersed. Species with fruit that contain pulp or aril were considered animal-dispersed, while those with fruits or seeds adapted for flight were considered wind-dispersed. For the details of the dispersal syndrome classification of only tree species, see the Supplemental Material.

For all analyses on the taxonomic and dispersal groups, we only included groups that had at least 5 species. We did not use a lower threshold on number of records for inclusion of a species, as species contributions to group-wide phenological dynamics are weighted by number of seeds in the analyses below.

### Weather data

We estimated monthly precipitation and minimum temperature at the plot level for each study site. Because local weather station data contained many missing observations, we used remotely sensed data. We used a ten-day precipitation time series estimated on a 0.05° grid by (Funk et al. 2014) using both remote and locally-sensed data. We used ECMWF/ERA-Interim reanalysis 4-hr temperature data at 2 m height estimated on a N128 Gaussian (~2°) grid (European Centre for Medium-Range Weather Forecasts 2009) and calculated daily minimum temperatures and then monthly values.

To estimate the rough pattern of wind seasonality in the study region, we used weather station data. For Cocha Cashu, we calculated average monthly wind speed from a station 150 km away, within 100 m elevation of Cocha Cashu, for the years 2004–2009 (http://atrium.andesamazon.org/meteo_station_display_info.php?id=12). For Yasuní, we used a weather station (http://www.serviciometeorologico.gob.ec/biblioteca/) 115 km away within 70 m elevation of the Yasuní plot for the years 2005-2012.

#### Statistical analysis

##### Wavelet transformation of seed rain data

For each species, we summed seed rain for each time point across all traps in the plot, resulting in a single time series for each species. Species seed abundance distributions were extremely right-skewed, with a few small-seeded species making up most seed counts. For example, the top five species make up 78% of seeds at Yasuní (2 Melastome species, 2 *Cecropia* species, and *Peperomia macrostachya*, all small-seeded), and 90% of seeds at Cocha Cashu (all five were small-seeded *Ficus*). To capture broader community patterns, rather than allowing patterns to be dominated by the few most abundant species, we transformed the counts to reduce skew (Vasseur et al. 2005, Lasky et al. 2016), specifically a natural log transformation of the counts + 1. To explore the sensitivity of our analyses to transformation, we also conducted all analyses in parallel without any transformation, however we focus our discussion on results from ln(counts + 1). In a subset of our analyses, we also explored how removing rarer species influenced our results (see Supplemental Material, also for a justification of using 1 as opposed to another constant). Note we did not address within-plot intraspecific variation in phenology as we were primarily interested in population- and community-wide phenology and its potential relationship to seed dispersers (that move across the plots) and temporal climate fluctuations (that are limited within the plot).

We then applied a continuous Morlet wavelet transformation to each species’ time series (either log transformed, or untransformed, greater detail in the Online Supplement). To characterize individual species’ phenology in relation to the community or group of species, we calculated the wavelet modulus ratio (WMR). WMR quantifies the relationship between the variation in the aggregate community-wide reproduction (numerator of Eqn. S3) relative to variation in species-level reproduction (Eqn. S3, Figure 1). When species seed rain dynamics through time cancel each other out, aggregate variation is zero (declining seed rain is balanced by increasing seed rain). Thus at zero, the WMR indicates compensation: all species-level dynamics are compensated so that community level reproduction is constant (Figure 1). At unity, the WMR signifies complete phenological synchrony among the species, as species-level phenological dynamics are completely reflected at the community level.

To answer Questions 1 and 2 above, we calculated the whole-community WMR for all species in Yasuní (0.10 to 8.5 yr periods) and Cocha Cashu (0.08 to 4.2 yr periods). The periods differed between sites because of the frequency of trap collection and duration. Trap observation dates were regularized to a constant interval following (Keitt 2014), and the minimum periods were twice the median sampling interval, limiting artifacts associated with a few days difference in intervals. The maximum scales were calculated as half the total duration of the study period at each site. We tested statistical significance of whole-community WMR using bootstrapping where each species had the phase of their wavelet at a given scale randomly shifted (Keitt 2008). This shift in phase has the effect of shifting where a wave is located in time, resulting in random patterns of among-species phenology. All analyses were run in R (v 3.3.2). WMR was calculated using the package ‘mvcwt’ (Keitt 2014, https://github.com/thk686/mvcwt).

##### Taxonomic and seed dispersal groups

To investigate if species that are closely related share similar phenological niches or partition phenological niches (Question 3 above), we focused on taxonomic groups. Our analyses were done at the family-level to ensure sufficient sample size. Confamilials often share characteristics making them likely to exhibit evolutionary niche conservation or character displacement. Similarly, groups with shared dispersal mechanisms might be more likely to exhibit non-random phenology (Question 3) so we separately grouped species based on their dispersal syndromes.

For these grouped analyses (family or dispersal syndrome), we first calculated WMR (as with the whole-community analyses above) for each taxonomic or dispersal group averaged across the time series. We excluded WMR values within one scale unit of the beginning and end of time series to reduce boundary artifacts (i.e. the cone of influence). To test the hypothesis that species within a group exhibited more synchronous seed fall or compensatory seed fall than the greater community, we generated a null distribution of each group’s WMR using permutations. We permuted species labels over the entire community while maintaining the number of species in each group, calculated WMR for the members of the permuted group, and then repeated this permutation 1000 times. If the observed WMR of a group averaged across time points was above the 97.5th percentile of the permutation-based null distribution, we considered it as nominally significant synchrony, if the observed WMR of a group averaged across time points was below the 2.5th percentile of the null distribution, we considered it as nominally significant compensation. We calculated two-tailed p-values from permutations and implemented false discovery rate (FDR) control across the multiple hypothesis tests using the method of (Benjamini and Yekutieli 2001).

##### Climatic association with synchrony vs compensatory dynamics

To determine whether climatic fluctuations might influence phenology among community members (Question 4 above), we investigated the association of local temperature and precipitation on whole community WMR calculated above. We calculated monthly average climate data and we aggregated seed rain data to monthly average seed counts. We then calculated WMR for Yasuní (2–70 mo scales) and Cocha Cashu (2–50 mo scales). We used wavelet transformation of the climate variables at the specific scales so that we could calculate their relationship with community WMR. Specifically, at each scale, we calculated the Pearson correlation coefficient between the community WMR and the wavelet-transformed minimum temperature or precipitation. Null distributions for correlations were generated by permuting the starting point of the wavelet transformed climate time series while maintaining periodic boundaries (i.e., adding climate values from before the randomly chosen staring points to the end of the permuted climate series) and calculating the Pearson correlation between the randomized climate wavelet and the WMR (n=1000, but with only 70 or 50 unique possible values for Yasuní and Cocha Cashu, respectively, and ignoring observations within one period of the boundary). The wavelet transform of climate was done in the R package “WaveletComp” (Rösch and Schmidbauer 2016).

## RESULTS

### Community-wide phenology

At the ever-wet site Yasuní we found a general trend of strong whole-community synchrony in transformed seed counts at scales of less than ~50 days, while larger sub-annual periods were weaker (significant at FDR = 0.05, Figure 2A). At the annual scale we also found significant synchrony for most of the study, and we found significant synchrony at scales greater than ~2 yrs, strongest at ~3.85 and 7 yrs (FDR = 0.05). These patterns were largely consistent (stationary) across the time period of the study, especially the pattern of strong synchrony at ~50 days. However, there was a weakening of annual-scale synchrony from 2012 to 2015. For untransformed seed counts, we found significant synchrony only at ~50 days and ~4 yrs (FDR = 0.05, Figure S2).

**Figure 2:**
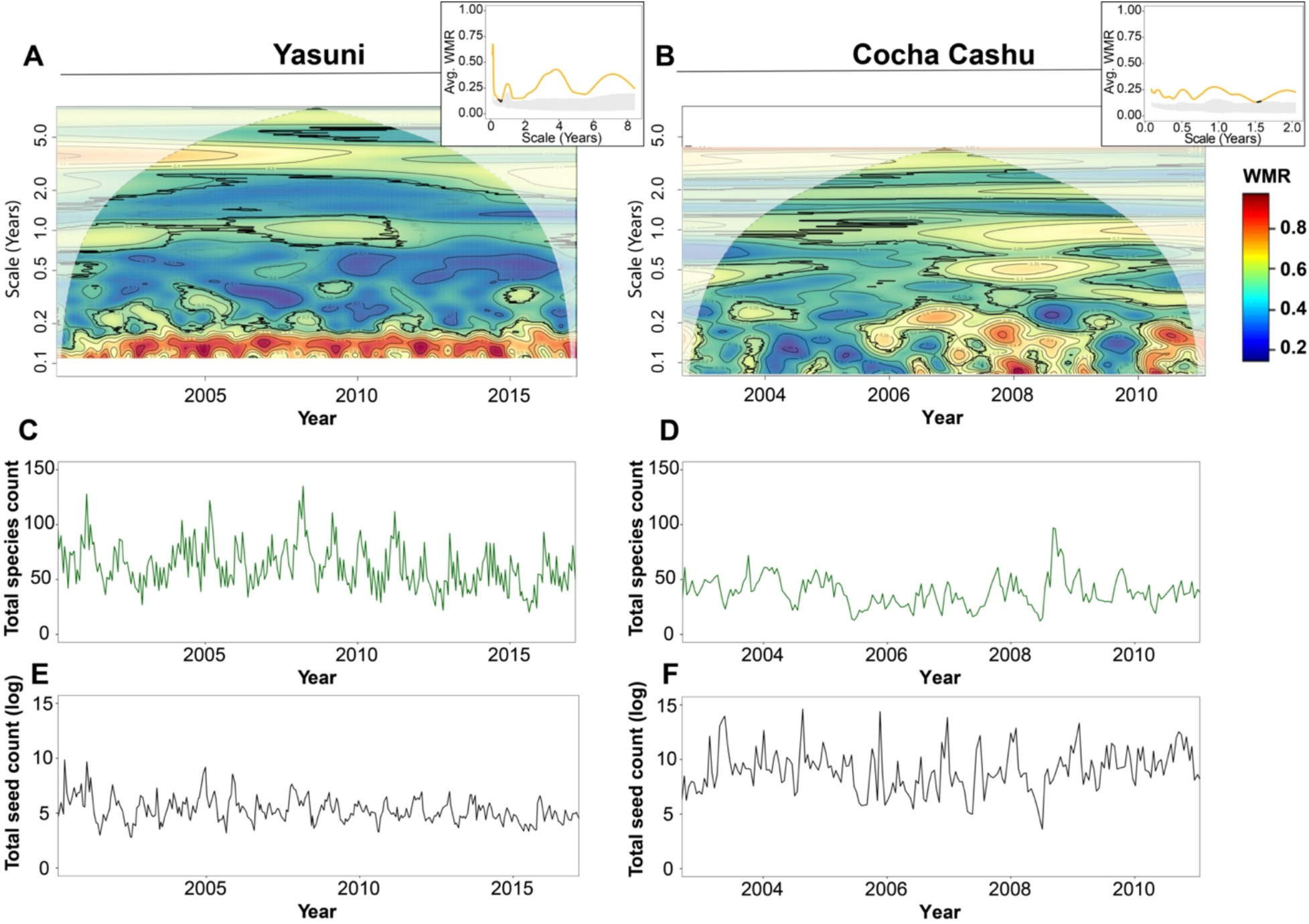
The whole-community wavelet modulus ratio (WMR) of log-transformed seed counts (A,B) at Yasuní (1059 species) and Cocha Cashu (654 species), and time series of the total species in traps (C,D) and total estimated seeds in traps (E,F, natural log) in each sampling period. In (A,B) red indicates high WMR while blue indicates low WMR. When WMR is greater than the null expectation this is significant synchrony, and when WMR is lower than expected we term this significant compensation. Insets (in A and B) show WMR averaged across the time series (gray ribbon is null distribution, the line indicates observed average WMR with yellow being scales where WMR is significantly different than the null) The thin dashed contour lines bound the points in time and scale (years) when the WMR was nominally significant (p<0.05) based on bootstrapping, while thick black lines bound regions significant with a false discovery rate (FDR) = 0.05. Nearly all significant regions in the plot are high WMR (yellow to red). The cone of influence (white shading, A,D) marks the regions where the wavelet transforms are affected by the boundaries of the sampling period. Note the null distribution of WMR values changes with scales. Log transformed counts are analyzed here. See Figure S2 for untransformed.

In contrast to Yasuní, at the seasonally dry Cocha Cashu site we found weaker whole-community synchrony in transformed counts at the sub-annual time scales before 2006, but for 2006 into early 2008 we found significant community-wide synchrony across a wide range of temporal scales (FDR = 0.05, Figure 2B). Additionally, there was consistent significant synchrony at the ~1 year and ~2 year, and >3 year scales across the duration of the study, indicating some shared annual, biannual, and multi-year dynamics among species (FDR = 0.05). However, for untransformed seed counts we found almost no evidence of non-random WMR patterns at Cocha Cashu (Figure S2).

### Phenology among confamilials

At Yasuní among the 28 families analyzed, we found that species of some families exhibited significant compensatory dynamics of transformed seed rain at sub-annual timescales (Figure 3A). Note that the null model for confamilials and dispersal groups was the community background, so significance indicates a stronger pattern than the rest of the community. In particular, species in the Annonaceae, Euphorbiaceae, Malpighiaceae, Myristicaceae, and Urticaceae families exhibited significant compensatory dynamics at different timescales up to 8 months (FDR = 0.05). That is, species that declined in reproduction over a few months tended to be replaced by other species in the same family increasing in reproduction over that timescale. Additionally, we found significant compensatory dynamics at the longer time scales (e.g. ~2.9 yrs) for Annonaceae. By contrast, at Cocha Cashu among the 27 studied families, some families exhibited synchrony and other families compensation, compared to the rest of the community. That is, confamilial species often showed significant synchrony or they showed significant compensation, especially at sub-annual time scales (Figure 3B). For untransformed data, we found only significant compensatory patterns within families, some of which (Annonaceae, Menispermaceae, Malpighiaceae) also were significantly compensatory in the analyses of transformed counts (FDR = 0.05, Figure S3).

**Figure 3:**
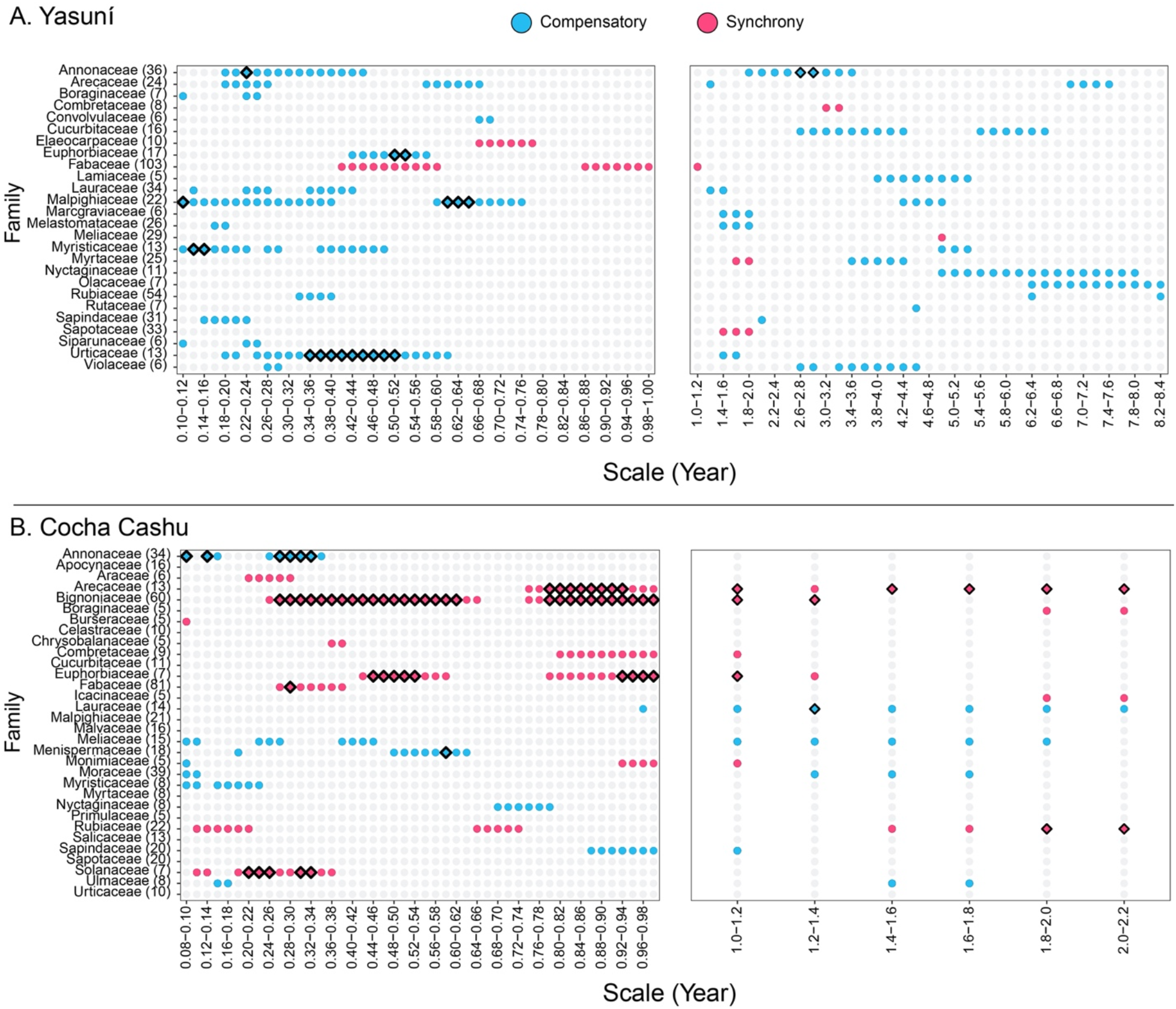
The averaged wavelet modulus ratio of families at Yasuní (A) and Cocha Cashu (B) at the sub-annual (left) and interannual (right) scales compared to a community-wide null. The number in parenthesis represents the number of species analyzed within the family. Colored points represent nominally significant (p<0.05) synchronous (red) or compensatory (blue) dynamics at the time scale. Thick borders around the points indicate significant points at false discovery rate (FDR) = 0.05. Log transformed counts are presented here. See Figure S3 for untransformed.

We found some consistency of family patterns across sites. Annonaceae at Cocha Cashu exhibited significant compensatory dynamics at sub-annual timescales (1–4 mos) similar to at Yasuní (transformed counts). Additionally, Myristicaceae at both sites showed nominally significant compensatory dynamics at 1–3 mo scales (though these were not significant after FDR control at Cocha Cashu). By contrast, Fabaceae species showed nominally significant synchrony for sub-annual to annual time scales at both sites (though these were not significant after FDR control at Yasuní, transformed counts).

To help illustrate the patterns identified, we present one family with synchrony among species and one family with compensatory dynamics among species (again compared to the other species in the community). At Cocha Cashu, Bignoniaceae shows strong synchrony at ~0.5 and 1 year scales, which corresponds to a strong twice yearly peak in seed fall of multiple species and many times of year with no seeds at all (Figure 4A,B). At Yasuní, Urticaceae showed significant compensatory dynamics at scales of ~0.34–0.52 years, which corresponds to a family-wide pattern where multiple species are typically releasing seed at any given point in time, and while some annual synchrony was evident, different species were often reproductive at different times of year so that seed counts never reached zero (Figure 4C,D, also see WMR figures for all families in Supplemental Material).

**Figure 4.**
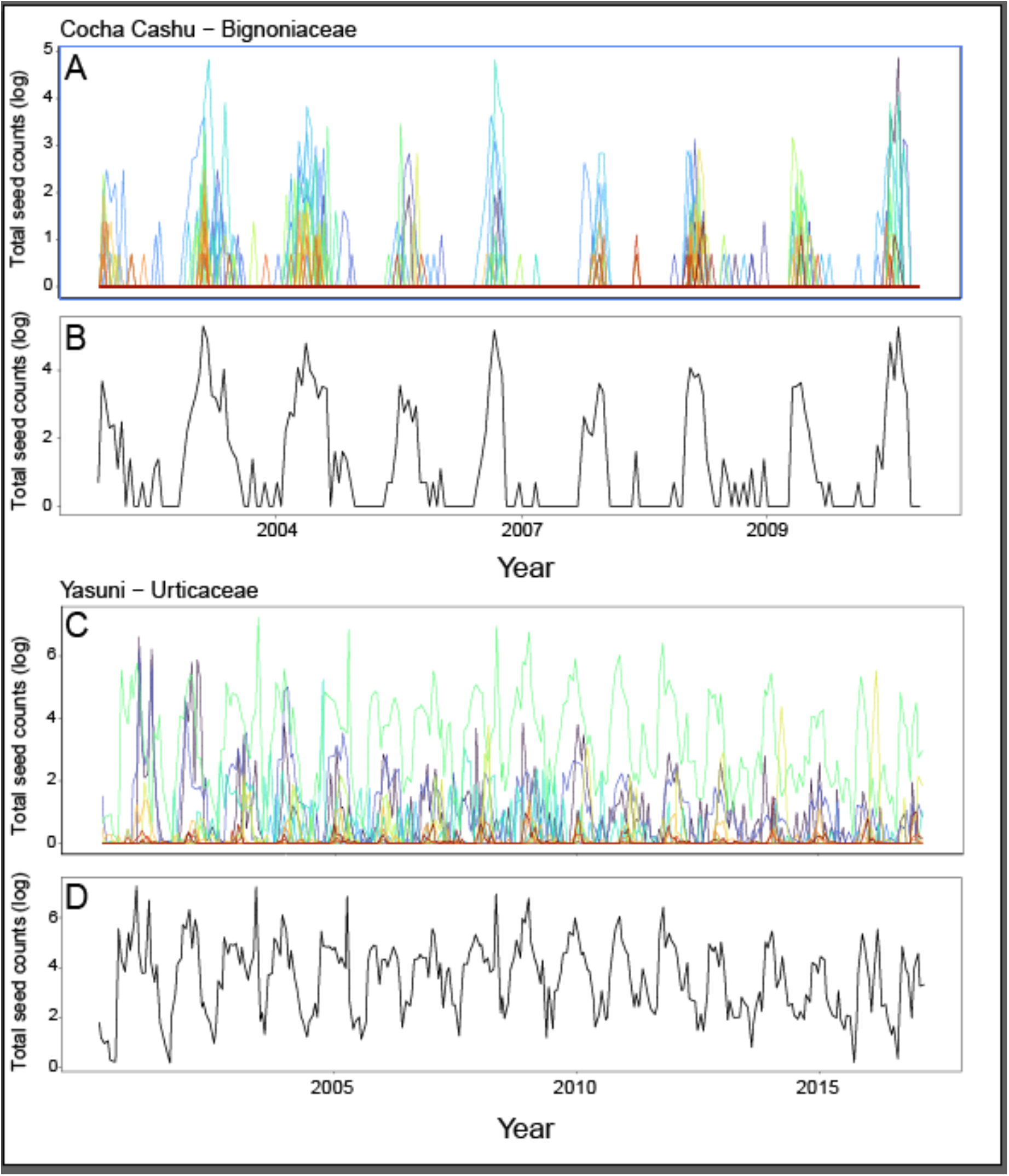
Two families at each site with distinct patterns of phenology, with each species’ time series of estimated seed counts in traps shown as a different colored line. Bignoniaceae (A-B) at Cocha Cashu (60 species) shows synchrony especially at ~1 year scales, with multiple species rising and falling in reproduction together in concert at these scales, and many times where no species are releasing seed. By contrast, Urticaeae (C-D) at Yasuní (13 species) shows significant compensatory dynamics compared the community-wide null at ~0.34–0.52 year scales. Although there is some synchrony, the most abundant Urticaeae species peak at different parts of the year, and even as some species decline in reproduction over these scales, others replace them, so that there is always substantial reproduction by some members of the family (D).

### Phenology among species sharing dispersal modes

Among the species with shared dispersal syndrome, we found significant synchrony in log transformed counts at multiple scales at the ever-wet Yasuní (Figure 5C & 5D). When considering all growth forms, we found that animal-dispersed species exhibited significant synchrony at ~4 month scales but no significant compensatory dynamics (FDR = 0.05). For wind-dispersed species, we found nominally significant (p < 0.05) synchrony at ~6 month scales that did not pass FDR control. For only trees, we did not find significantly non-random phenology for groups of species with similar size fruits or similar abiotic dispersal mechanisms (though some were nominally significant p < 0.05, Figure S5). Untransformed seed counts generally did not show significantly different WMR within dispersal groups at Yasuní (Figures S4 and S6).

**Figure 5.**
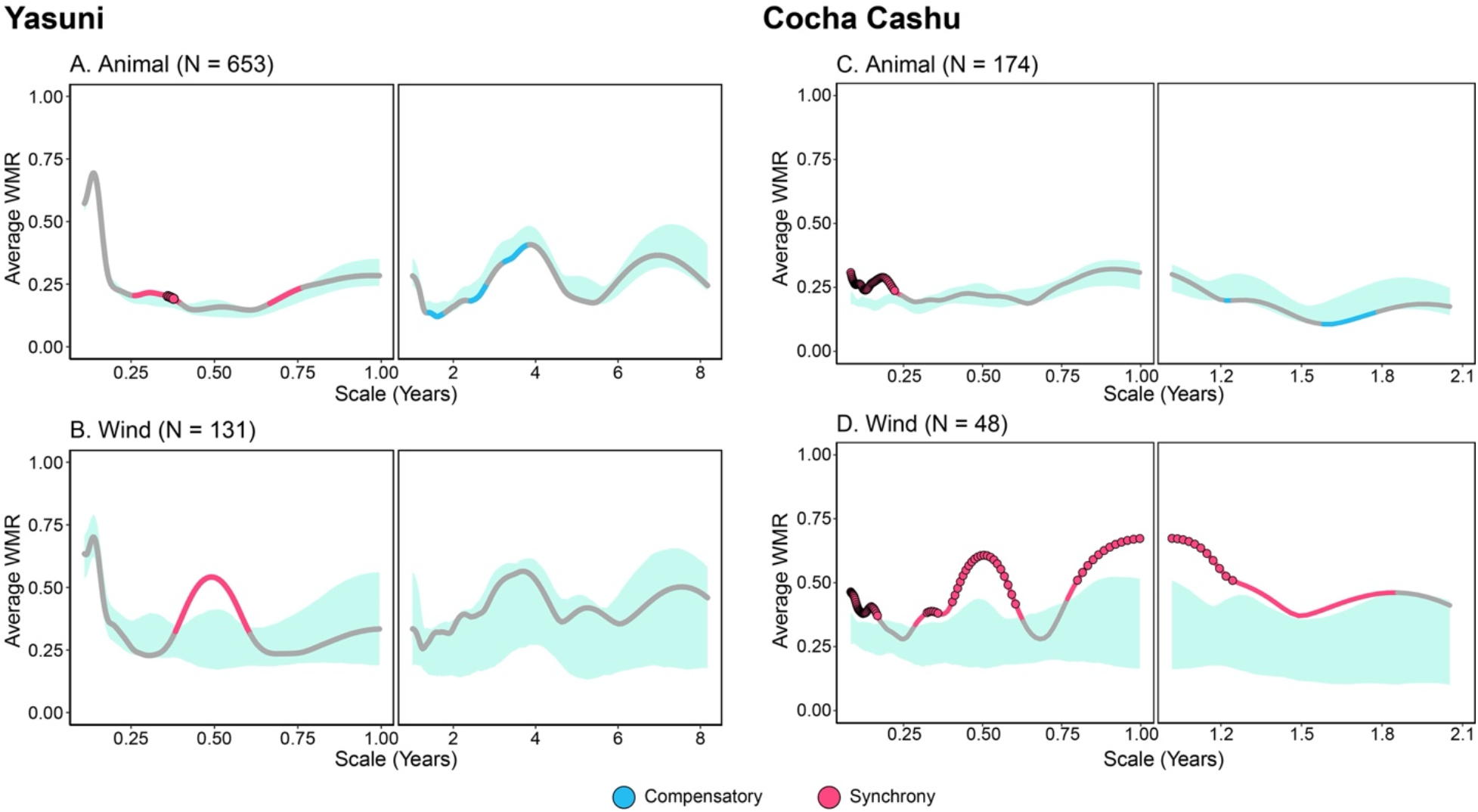
The averaged wavelet modulus ratio for plant species of all growth forms based on dispersal syndrome (thick lines). The number in parentheses represents the number of species within the animal and wind-dispersed groups. The light blue ribbon represents the 2.5–97.5^th^ percentiles of the null-distribution generated through bootstrapping. Any points that lie above the ribbon were considered nominally significant and synchronous while any points below the ribbon were considered nominally significant and compensatory dynamics (p<0.05). Enlarged points indicate significant at FDR = 0.05.

At seasonally dry Cocha Cashu, we found significant synchrony among log transformed seed counts for dispersal groups of all growth forms (Figure 5). Wind-dispersed species exhibited strong and significant synchrony at ~1, ~6, and ~12 month timescales (FDR = 0.05, Figure S9). Wind variation also showed a peak in variability at ~12 month scales (and ~6 month, depending on the metric, Figure S7). Animal-dispersed species showed significant synchrony at ~1–3 month scales (FDR = 0.05). For only tree species, we did not find non-random phenology for species with similar dispersal syndromes, except for ~1–2 month synchrony for large vertebrate-dispersed species (FDR = 0.05, Figure S8). Untransformed seed counts within dispersal groups were not significantly different from the null, with the exception of animal-dispersed species of all growth forms at ~1.7 yr scales (Figures S10–11).

### Temperature, precipitation, and community-wide phenology

There was a significant relationship between temperature (but not precipitation) and community-wide WMR for log transformed seed counts (FDR = 0.05, Figure S13–14). Yasuní had a positive temperature–WMR relationship most strongly at ~1.5 and 2 yr scales, indicating periodic increases in among-species seed rain synchrony with warming temperatures. At the seasonally dry Cocha Cashu at ~1.3 yr scales, there was a significant positive correlation between the WMR and minimum temperature, similar to Yasuní. Unexpectedly, there was no increase in synchrony at Cocha Cashu during wet or dry periods. There were some similar patterns observed in the untransformed counts’ WMR– climate correlations (Figure S15-16).

## DISCUSSION

Ecological communities harbor extensive phenological diversity among member species, especially in tropical wet forests (Frankie et al. 1974). This phenological diversity might help explain species coexistence within communities if phenology is a key axis of niche partitioning among species. Despite this great phenological diversity, there are well-known examples of strong synchrony in tropical wet forests that lack strong seasonality (Ashton et al. 1988). However, temporal traits (e.g. aspects of phenology) remain lesser-known dimensions of diversity in ecological communities.

We found evidence that Amazonian plant communities frequently exhibit significant reproductive synchrony at certain time scales at the whole-community level, suggesting that shared responses to environment or positive interactions among species structure community phenology. However, groups of species sharing dispersal mechanisms or related groups of species sometimes exhibited compensatory (anti-synchronous) and sometimes synchronous dynamics.

### Do communities exhibit synchronous or compensatory reproduction?

Overall, we found many cases of strong synchronous dynamics in seed rain at the whole-community level, where across the whole community species tended to rise and fall in seed rain in concert. We found almost no significant whole-community compensatory dynamics. The synchrony we observed suggests whole-community dynamics are driven by shared responses to fluctuations in environmental (abiotic or biotic) conditions such as rainfall or frugivore abundance, or by positive heterospecific interactions such as enhanced frugivore attraction or natural enemy satiation (van Schaik et al. 1993). Note that in our study we were only able to observe patterns deviating from our permutation-based null. Processes that influence the shape (but not the phase, which we randomized to generate our null, see Figure 2 A, B insets) of the phenology of the species pool are outside our scope of detection (Case et al. 1983).

The ecological forces shaping phenology of fruiting and seed release are also tied to other developmental stages. Flowering, leaf emergence and senescence, and seed germination all can interact with abiotic and biotic conditions. Future research will benefit from an integrative understanding of how full life-cycle phenology interacts with environment (Borchert 1996).

### Are compensatory and synchronous dynamics scale-specific or non-stationary?

At both sites, synchronous seed fall was strongest and most consistent at scales of approximately 1–2 mos, 1 yr, and >3 yrs. The fastest scale of synchrony at ever-wet Yasuní (~1.5-2 mos) may represent shared and rapid community responses to relatively brief cloudless periods of high radiation alternating with cloudy periods, similar to what was observed in response to rainfall in a seasonally dry forest (Lasky et al. 2016). However, we did not find a correlation between temperature or rainfall and WMR at the ~1 mo scale, and inconsistent correlations at larger time scales. At both sites, the yearly and super-annual patterns of synchrony were fairly consistent throughout the study. Clear annual peaks were visible even in the raw species or seed counts over time, highlighting the strength of synchrony at this scale, despite the presence of many reproductive species year-round (Figure 2 C, D, E, F).

At seasonally dry Cocha Cashu, we observed strong non-stationarity, in particular a shift from approximately random to synchronous phenology, across all time scales <1 yr, with a peak in WMR in 2008 (Figure 2B). The ecological explanation for this shift is unclear, but in 2008 there was the lowest nadir in species and total seed counts followed shortly by the highest peak in species number (Figure 2 D, F). The decrease followed by a high peak in reproduction might indicate multiple species were accumulating resources synchronously (reducing reproduction) in order to subsequently invest a large amount in reproduction, akin to bursts of reproduction in masting species (Janzen 1974, Ashton et al. 1988).

### Is evidence for phenological partitioning strongest among ecologically similar species?

In contrast to our findings at the whole-community level, we often found significant compensatory dynamics within focal groups of species (confamilials and species with similar animal seed dispersers), regardless of whether or not we transformed seed counts. Whole-community patterns may obscure phenological niche partitioning that occurs within groups of closely interacting species. This hypothesis is consistent with findings of ecological character displacement among closely related species in a wide variety of taxa (Robinson and Wilson 1994, Dayan and Simberloff 2005), including in time series of phytoplankton populations (Barraquand et al. 2018). Previous studies that have shown evidence for such partitioning have been largely focused on groups of closely related species hypothesized to be closely interacting due to shared pollinators or mutualists (Gentry 1974, Ashton et al. 1988, Botes et al. 2008). Future efforts might also identify these species based on scale-specific, non-random phenological patterns instead of relying on prior ecological or phenotypic knowledge. Annonaceae species showed similar patterns of compensatory seed fall for within-year timescales at both sites, perhaps indicating consistency in phenological niche partitioning among species in this family. Many Annonaceae species have a similar fruit morphology of fleshy stipitate monocarps and animal dispersal, such that phenological niche partitioning may promote coexistence.

It is challenging with our approach to determine the specific mechanisms behind any phenological niche partitioning. An example misleading scenario is where species share dispersal mechanisms but also other traits, of which the latter lead to resource competition that is ameliorated by phenological niche partitioning. To demonstrate niche partitioning with respect to animal dispersers at our sites requires additional evidence. For example, (Botes et al. 2008) showed how species that deposit pollen on the same location on pollinators’ bodies (suggesting competition or interference) exhibited compensatory flowering, but species depositing pollen on different body locations did not.

### Synchrony among species sharing dispersal mode or related species

We found that among all growth forms, wind-dispersed species exhibit strong synchrony of seed rain compared to the whole community, particularly at time scales indicative of shared abiotic niches. At both sites we found some degree of ~6 mo synchrony among wind-dispersed species (as well as the wind-dispersed family Bignonicaceae at Cocha Cashu), consistent with the twice-yearly peaks in wind speed observed at nearby weather stations (Figures S7, S9, S12). Near Cocha Cashu there is a strong peak in wind speed in September (Figure S7), also consistent with the synchrony at this site observed among wind-dispersed species at ~1 yr periods (Figures 5 & S9). Near Yasuní the wind had a broad peak in average speed from September to November with a small peak in April (Figure S12). The tendency for wind-dispersed tropical species to synchronously release seed during windy seasons has been reported in the literature (Frankie et al. 1974, Janzen 1974, Detto et al. 2018) and can be considered a positive control for our approach.

We found significantly greater synchrony for several speciose families at ~1 yr scales or sub-annual scales for Arecaceae, Euphorbiaceae, Fabaceae, and Solanaceae (in addition to aforementioned Bignoniaceae at Cocha Cashu). Given the dominance of climate variation at 1 year scales (Figures S14–15) the ~1 year synchrony in these families may be tied to shared responses to abiotic fluctuations, although there are no clear ecological similarities among these ecologically distinct families. Our finding may partly indicate greater power to detect patterns in more speciose groups.

### Community phenology and abiotic fluctuations

Gentry (1974) hypothesized that phenological diversity of communities was promoted by permissive abiotic conditions and longer growing seasons compared to seasonally harsh environments, where the range of potential phenologies is narrower. We saw mixed evidence for this hypothesis based on fluctuations in climate at our sites. At both sites, sub-annual WMR was only weakly (if significantly) associated with temperature. Super-annual WMR showed some strong associations with temperature, in particular at Yasuní were we found at >4.5 yr scales higher synchrony in warmer periods. Similarly, at Barro Colorado Island in Panama, community-wide peaks in seed rain occur during ENSO events (Detto et al. 2018), presumably as trees had shared (synchronous) responses to increased light during these dry and warm periods (Wright and Calderón 2006). However, we also found lower WMR in warmer periods at ~3.3 yr scales at Yasuní, leaving our conclusions unclear.

We did not observe differences between ever-wet Yasuní and seasonally dry Cocha Cashu that can easily be explained by the sites’ precipitation seasonality. Even though Yasuní may represent the most climatically favorable site, the whole community still showed strong synchrony. Furthermore, there was no link between precipitation and the strength of synchrony at seasonally dry Cocha Cashu (Figure 6). While we found temporal associations with climate, we had too few sites to ask whether spatial climatic differences are associated with whole-community differences in synchrony and compensation. Also note that the larger area covered by traps at Yasuní and the longer time series may have increased the precision of our estimates of population phenology and our power to detect non-random patterns, compared to Cocha Cashu.

### Conclusion

Here we showed how whole-community phenology in diverse plant communities is largely characterized by synchrony. However, we also found that groups of related or ecologically similar species often show compensatory patterns of seed rain, indicating potential phenological axes of niche partitioning that might promote species coexistence. Our results highlight the scale-specific and sometimes non-stationary characteristics of community phenology that may arise from the multiple opposing drivers of phenology. These changing patterns across temporal scales are similar to those observed in trait based tests of community assembly across spatial scales, where authors have found trait diversity patterns can reverse across scales indicating potentially opposing community assembly forces acting at different scales (Swenson and Enquist 2009). Flexible multi-scale analyses may reveal evidence of scale-specific niche partitioning and environmental filtering and suggest the mechanisms that structure communities.

## Acknowledgements

This manuscript benefited from comments of Tomás Carlo, Kathryn Cottingham, and two anonymous reviewers. Work at Yasuní was supported by funding to NCG and collaborators from the Andrew W. Mellon Foundation, Natural Environment Research Council (GR9/04037), British Airways, Department of Botany, Natural History Museum, and the National Science Foundation (DEB-0614525, DEB-1122634, DEB-1754632, DEB-1754668). We thank the Ecuadorian Ministerio del Ambiente for permission to work in Yasuní National Park (under N° 014-2019-IC-PNY-DPAO/AVS, N° 012-2018-IC-PNY-DPAO/AVS, N° 008-2017-IC-PNY-DPAO/AVS, N°. 012–2016-IC-FAU-FLO-DPAO-PNY, N°. 014-2015-FLO-MAE-DPAO-PNY, and earlier permits). We very gratefully thank Milton Zambrano for collecting most of the Yasuní trap data from 2002–2017. We also thank Viveca Persson for help initiating the censuses in 2000–2002, with assistance from Zornitza Aguilar, Paola Barriga and Matt Priest, and Gorky Villa, Alvaro Perez and Pablo Alvia for help identifying species. Data collection at Cocha Cashu was supported by funding from the Andrew W. Mellon Foundation and the National Science Foundation (DEB-0742830). We thank the Peruvian authorities INRENA and SERNANP for permission to work in Manu National Park (under N° 020-CC-2008-INRENA-IANP, 05-CC-2009-SERNANP-PNM, 010-2010-SERNANP-JPNM, and earlier permits). More than 25 Peruvian undergraduate students assisted with data collection from 2002-11. Vishnu Viswanathan provided assistance digitizing weather records.

