## Supplemental Material for "Multi-scale phenological niches of seed fall in diverse Amazonian plant communities"

**Supplementary Material**

*Seed rain data*

Some species at Yasuní were not well separated in earlier years and thus these species were excluded from some analyses. For the family-level analyses, we censored Clusiaceae (due to issues with *Clusia* identification) and Solanaceae before 1/1/2007. For the family-level analyses of Moraceae, time series were censored before 1/1/2008 (due to issues with *Ficus* identification). Species without identification issues in these families were included in whole-community analyses.

At Yasuní, seeds and whole mature fruits were counted individually; fruit segments (such as capsule values) were aggregated and counted as the equivalent number of whole fruit. The number of seeds per fruit was counted directly from fresh specimens, our reference collection or photographs, or estimated from generic or familial data. These estimates of seeds per fruit were used to impute seed number from counted fruits.

For fruits collected at Cocha Cashu, fruit counts were converted to seed counts by multiplying by the average number of seeds per fruit for that species. Data on seeds per fruit were obtained from the literature (Alvarez-Buylla and Martinez-Ramos 1992, Gentry 1996, Kalko and Condon 1998, Stevenson et al. 2002, Russo 2003, Cornejo and Janovec 2010). For both sites, unidentified seeds not counted as specific morphospecies were excluded.

*Tree seed dispersal syndromes*

For trees with animal-dispersed seeds at Yasuní we classfied syndromes as follows: dispersed by terrestrial animals (25 species), or dispersed by canopy animals with small (< 2cm, 230 species), medium (2–5 cm long, 74 species), or large (> 5 cm, 12 species) seeds (groups are mutually exclusive). Abiotic-dispersed trees were split into ballistically dispersed (16 species) and wind dispersed (30) species.

For dispersal syndrome classifications at Cocha Cashu, Bagchi et al. (2018) previously estimated each species' proportional dispersal by members of seven dispersal groups, using information from published studies conducted in the Madre de Dios basin and other long-term Neotropical rain forest sites. The groups were: 1) large- and medium- bodied vertebrates (e.g. tapirs, spider monkeys, capuchins, guans, toucans, trumpeters, 52 species), 2) small bodied non-volant arboreal mammals (e.g. tamarins, night monkeys, kinkajous, 20 species), 3) small birds (e.g. manakins, cotingas and tanagers, 25 species), 4) bats (5 species), 6) wind dispersal (8 species), and other smaller categories/unknowns (Bagchi et al. 2018). We took these continuously-varying published estimates and had to cluster them into discrete groups to calculate dispersal-group specific WMR. Specifically, we performed k-means clustering to produce six mutually exclusive groups of species with similar dispersal modes.

*Wavelet analyses*

For each species, we summed seed rain for each time point across traps and then log-transformed the count + 1, resulting in a single time series for each species. We then applied the continuous wavelet transformation to each species' time series using the Morlet wavelet

$$W_k\,(a,\tau) \;=\; \frac{1}{\sqrt{a}}\int_{-\infty}^{\infty} x_k(t)\varphi\frac{t-\tau}{a}d\tau \qquad \text{(Equation 1).}$$

Here, the wavelet coefficient $W_x$ is the cross-correlation between species' $k$ seed rain time series, $x_k$(t), and the complex-valued Morlet wavelet $\varphi$,

$$\varphi(\tau) = \pi^{-\frac{1}{4}}\exp\,(2\pi i\tau - \tfrac{1}{2}\tau^2) \qquad \text{(Equation 2).}$$

A complex Morlet wavelet is a Gaussian-tapered complex sine wave, where the tapering allows one to capture localized patterns. The wavelet is stretched to different scales, $a$, such that the Gaussian taper occurs over different scales, and translated across the different points in time of the study, $\tau$.

When the counts $x_k$(t) in Eqn 1 are zero, the variation in wavelet function is not transmitted to the transformed wavelet coefficient $W_k\ (a, \tau)$. This is true for both raw untransformed counts, or log($x$ + 1) transformed counts. However, if a different constant is added to $x$ in the log transformation, the "baseline" count value (i.e. when truly zero seeds were observed) is no longer zero, and as a result the variation in the wavelet function $\varphi$ is transmitted to the wavelet coefficients, despite no variation in true seed rain. In this case, the only variation in the wavelet coefficient arises from the wavelet function $\varphi$, rather than the seed counts. This constraint justifies the use of log($x$ + 1) as opposed to some other constant instead of 1. Using other constant values results in wavelet variation transmitted to the transformed wavelet coefficients $W_k\ (a, \tau)$ even when there is constant zero seed rain. Most species have many zero counts, and as a result, artificial synchrony (high WMR) is introduced. See Figures S20-S23.

After each species' seed rain time series was wavelet transformed, we then sought to characterize each species' phenology in relation to the entire community of species, or taxonomic/dispersal group of species. To do so, we calculated the wavelet modulus ratio (WMR). This quantifies the relationship between the variation in the aggregate community-wide reproduction (numerator of Eqn. 3) relative to variation in species-level reproduction (denominator of Eqn. 3) at scale $a$ and centered on time $t$,

$$WMR(t, s) = \frac{\Lambda_{t,a}(|\sum_k W_k(\tau,a)|)}{\Lambda_{t,a}\ \sum_k |W_k(\tau,a)|} \qquad \text{(Equation 3)}$$

where $\Lambda_{t,a}(.) = \int_{-\infty}^{\infty} e^{-\frac{1}{2}\left(\frac{t-\tau}{a}\right)^2}(\cdot)d\tau$ and $|\cdot|$ represents the complex norm (Keitt 2008, Keitt 2014, Lasky et al. 2016) (Figure 1). When species seed rain dynamics through time perfectly cancel each other out, the sum in the numerator of Eqn 3 is equal to zero (declining seed rain is balanced by increasing seed rain). Thus at zero, the WMR indicates complete compensation or anti-synchrony: all species-level dynamics are compensated so that community level reproduction is constant. At unity, the WMR at the time period signifies complete phenological synchrony among the species, as species-level phenological dynamics are completely reflected at the community level.

**Figure S1.** Map showing locations of the Yasuní (YAS) and Cocha Cashu (CC) plots.

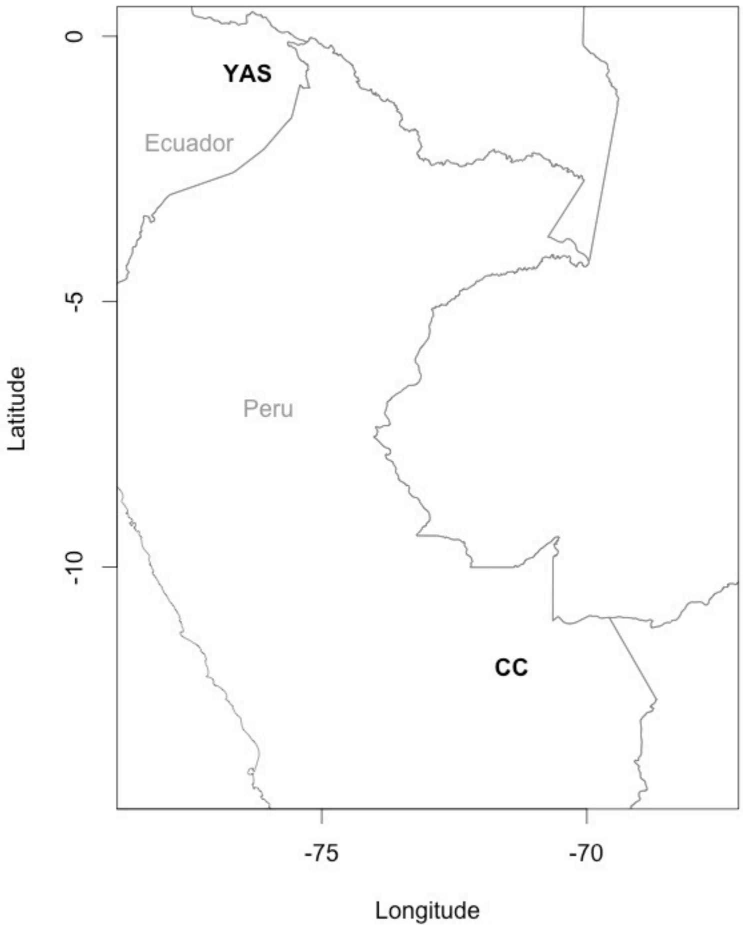

**Figure S2.** The whole community wavelet modulus ratio (WMR) of raw untransformed seed counts (A,B) at Yasuní (1059 species) and Cocha Cashu (654 species), and time-series of the total species in traps (C,D) and total estimated seeds in traps (E,F, natural log) in each sampling period. In (A,D) red indicates high WMR while blue indicates low WMR. When WMR is greater than the null expectation we refer to this as significant synchrony, and when WMR is lower than expected we term this significant compensatory dynamics. The thin dashed contour lines bound the points in time and scale (years) when the WMR was nominally significant (p<0.05) based on bootstrapping, while thick black lines bound regions significant with a false discovery rate (FDR) = 0.05. The cone of influence (white shading, A,D) marks the regions where the wavelet transforms are affected by the boundaries of the sampling period. Note the null distribution of WMR values changes with scales, and insets (in A and B) show averaged behavior across the time series (gray ribbon is null distribution, the line indicates observed average WMR with yellow being scales where WMR is significantly different than the null). See Figure 2 for log transformed WMR.

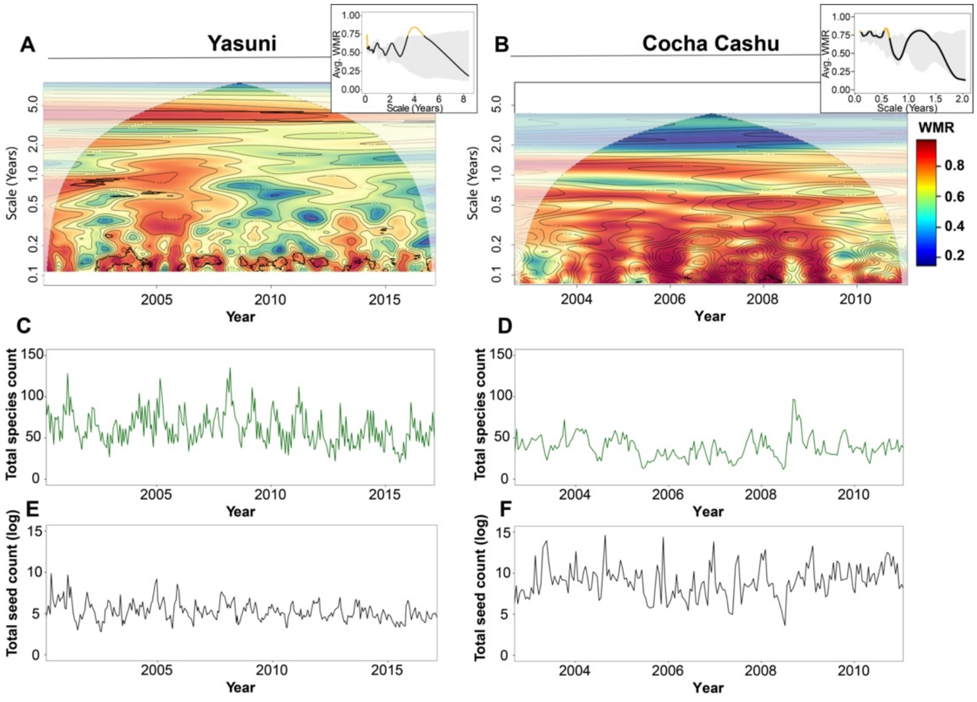

**Figure S3.** The averaged wavelet modulus ratio of families at Yasuní (A) and Cocha Cashu (B) for raw untransformed counts, at the sub-annual (left) and interannual (right) scales compared to a community-wide null. The number in parenthesis represents the number of species analyzed within the family. Colored points represent either nominally significant synchronous (red) or compensatory (blue) dynamics at the time scale ($p < 0.05$). Thick borders around the points indicate significant points at false discovery rate (FDR) = 0.05. See Figure 3 for log transformed.

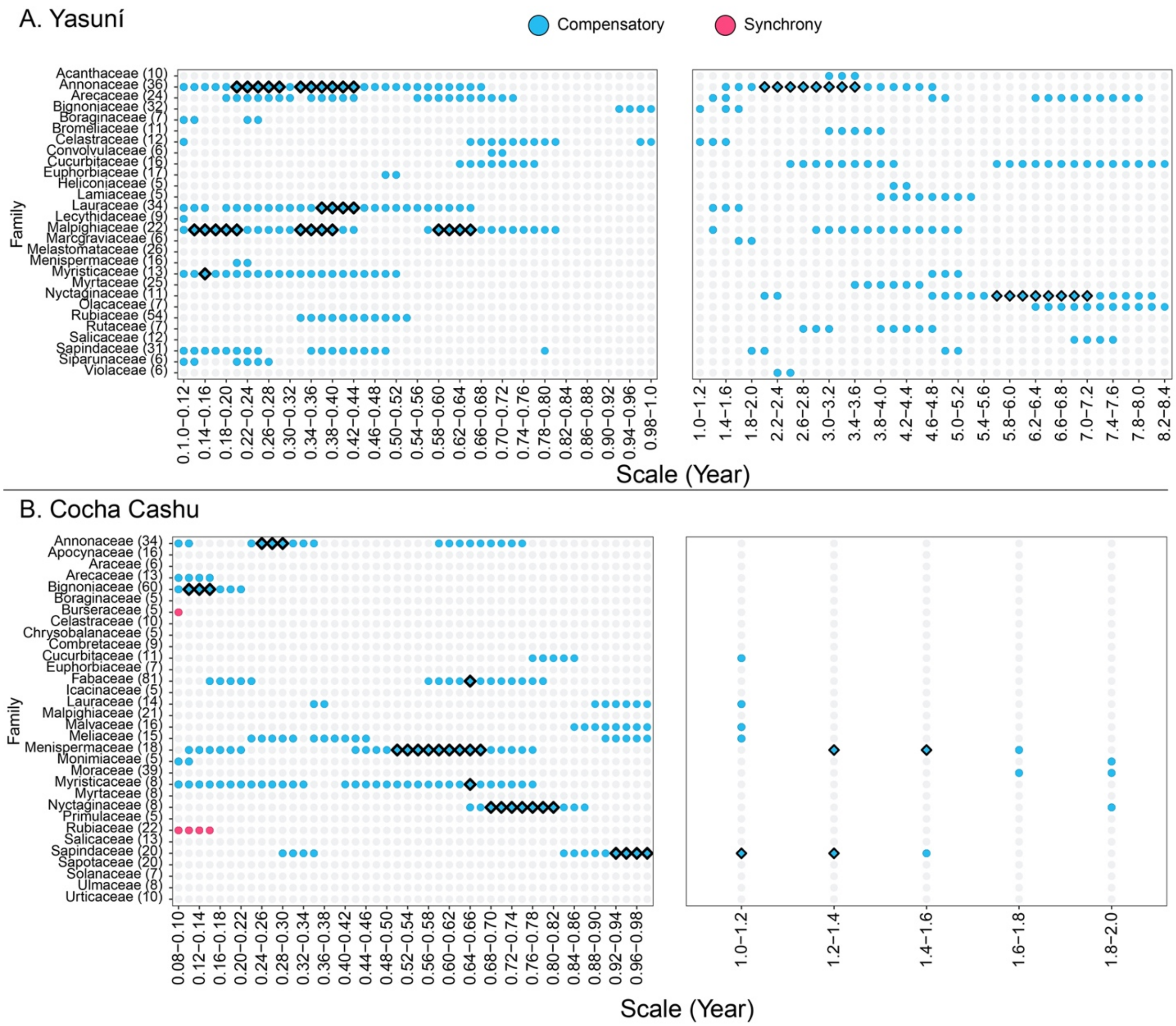

**Figure S4.** The averaged wavelet modulus ratio for plant species of all growth forms based on dispersal syndrome (thick lines), here showing raw untransformed counts for Yasuní. The number in parentheses represents the number of species within the animal and wind-dispersed groups. The light blue ribbon represents the 2.5-97.5th percentiles of the null-distribution generated through bootstrapping. Any points that lie above the ribbon were considered nominally significant and synchronous while any points below the ribbon indicated nominally significant, compensatory dynamics ($p<0.05$). For log transformed counts see Figure 5.

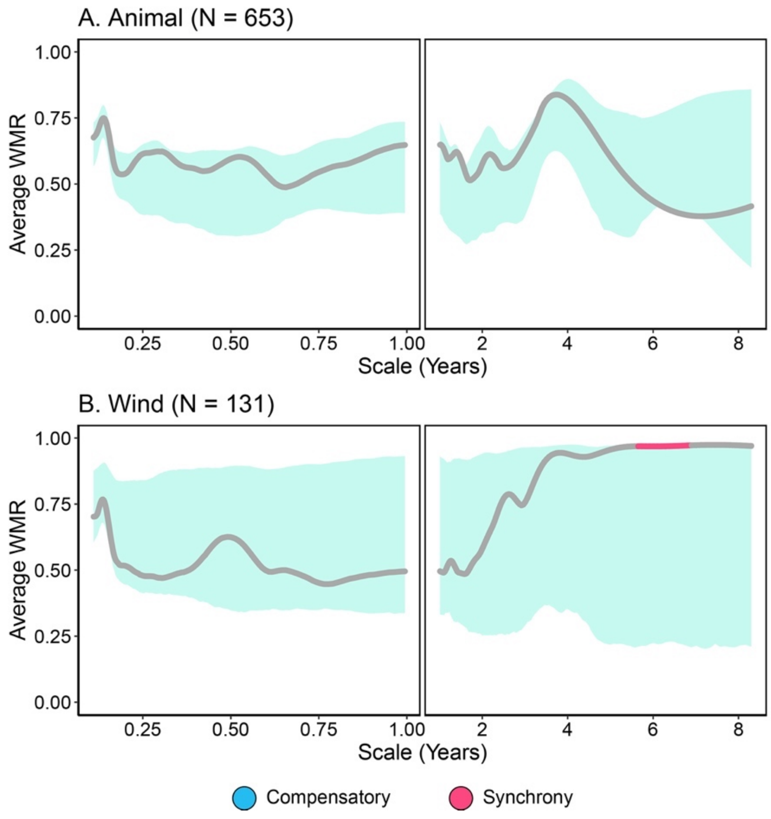

**Figure S5.** The averaged wavelet modulus ratio for tree species based on dispersal syndrome (thick lines), here showing log transformed counts for Yasuní. The number in parentheses represents the number of species within the groups. The light blue ribbon represents the 2.5-97.5th percentiles of the null-distribution generated through bootstrapping. Any points that lie above the ribbon were considered nominally significant and synchronous while any points below the ribbon indicated nominally significant, compensatory dynamics ($p<0.05$). For untransformed counts see Figure S6.

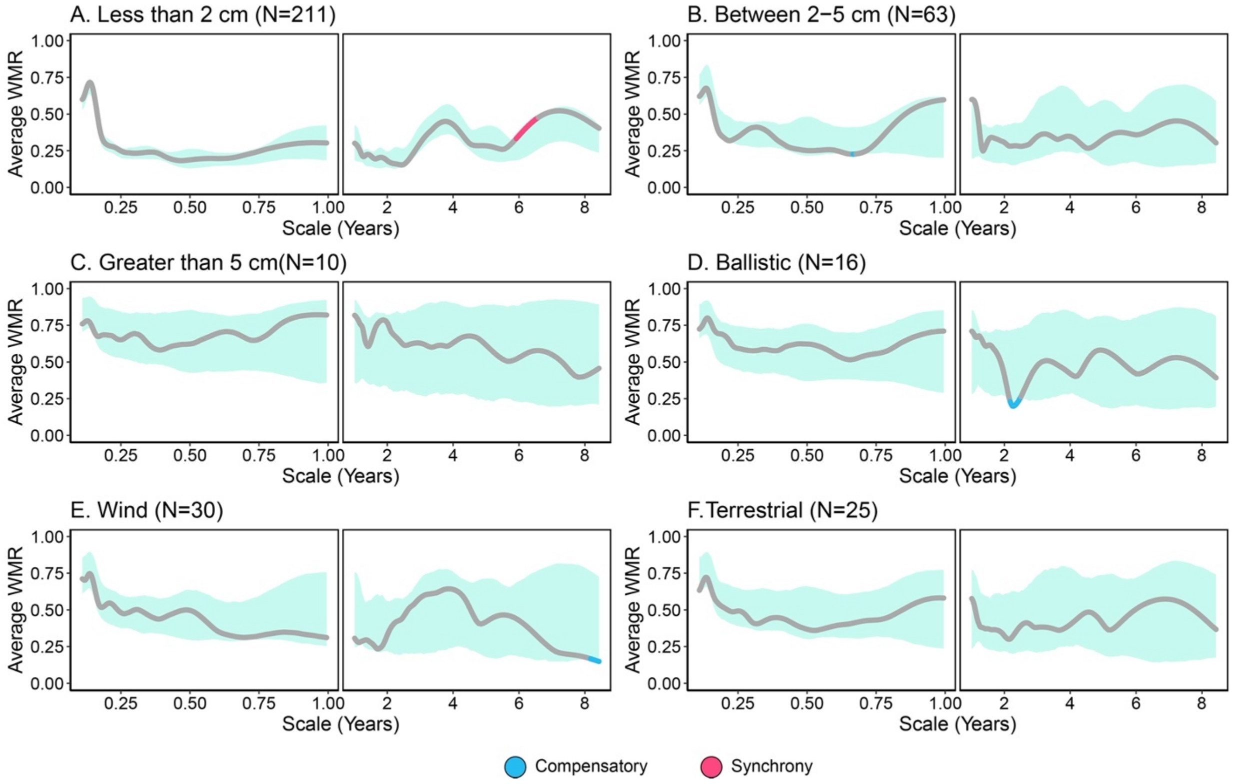

**Figure S6.** The averaged wavelet modulus ratio for tree species based on dispersal syndrome (thick lines), here showing raw untransformed counts for Yasuní. The number in parentheses represents the number of species within the groups. The light blue ribbon represents the 2.5-97.5th percentiles of the null-distribution generated through bootstrapping. Any points that lie above the ribbon were considered nominally significant and synchronous while any points below the ribbon indicated nominally significant, compensatory dynamics ($p<0.05$). For log transformed counts see Figure S5.

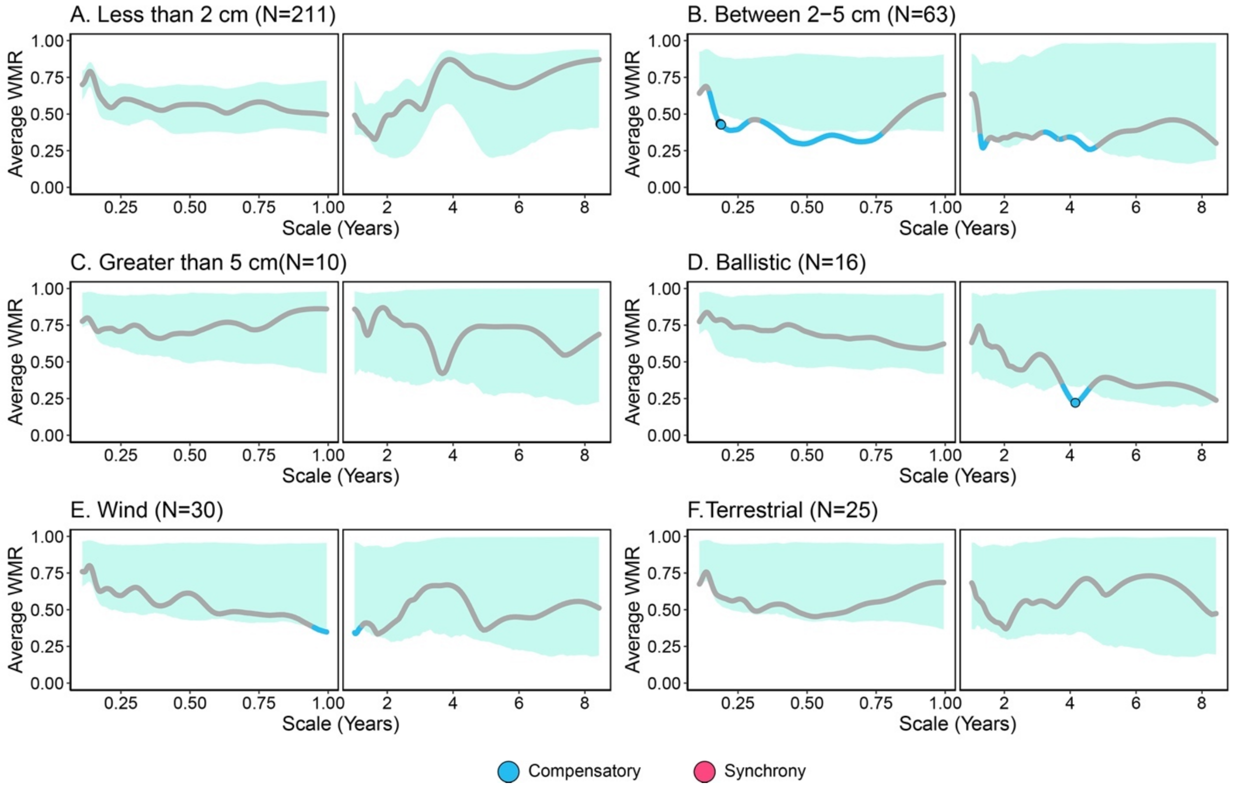

**Figure S7.** Wavelet power (top panel) and monthly averages (bottom panel) for average wind speed at a weather station near Cocha Cashu, 2004-2009.

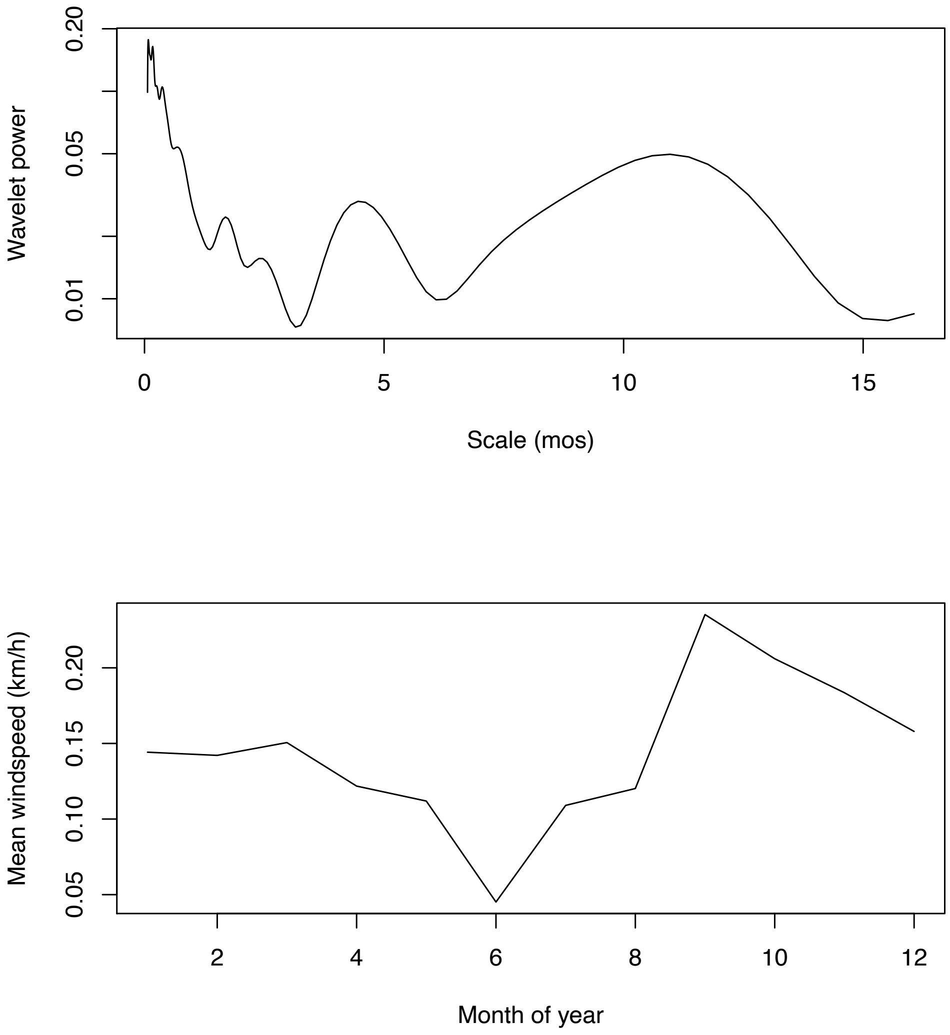

**Figure S8.** The averaged wavelet modulus ratio for tree species based on dispersal syndrome (thick lines), here showing log transformed counts for Cocha Cashu. The number in parentheses represents the number of species within the groups. The light blue ribbon represents the 2.5-97.5$^{th}$ percentiles of the null-distribution generated through bootstrapping. Any points that lie above the ribbon were considered nominally significant and synchronous while any points below the ribbon indicated nominally significant, compensatory dynamics ($p<0.05$). For log transformed counts see Figure S11.

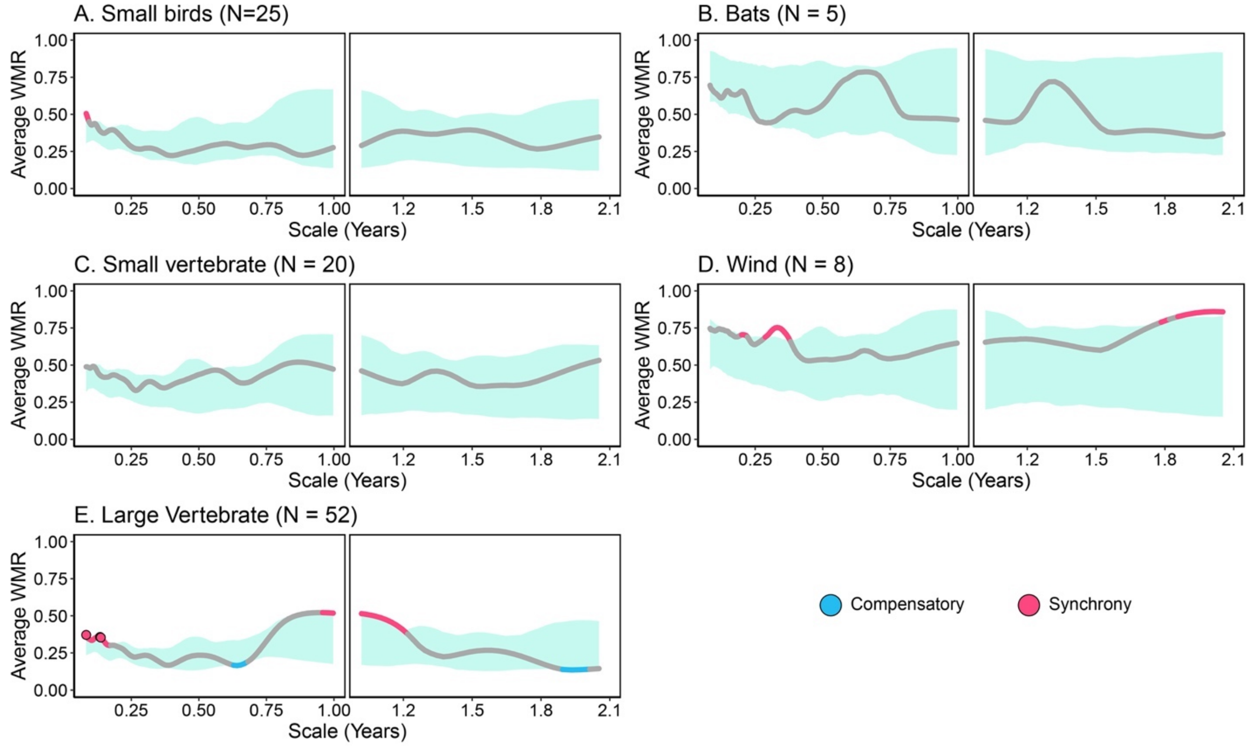

**Figure S9.** Time series of wind-dispersed seed counts (with a 6 week smooth for visualization) at Cocha Cashu, highlighting the twice yearly peaks and 6 month scale synchrony.

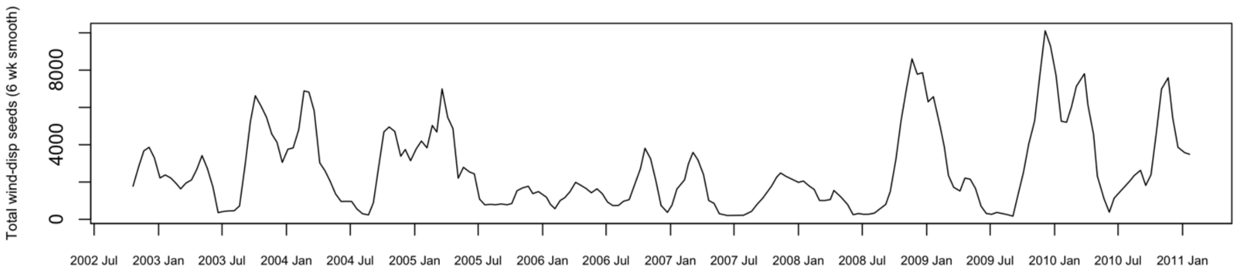

**Figure S10.** The averaged wavelet modulus ratio for plant species of all growth forms based on dispersal syndrome (thick lines), here showing raw untransformed counts for Cocha Cashu. The number in parentheses represents the number of species within the animal and wind-dispersed groups. The light blue ribbon represents the 2.5-97.5$^{th}$ percentiles of the null-distribution generated through bootstrapping. Any points that lie above the ribbon were considered nominally significant and synchronous while any points below the ribbon indicated nominally significant, compensatory dynamics ($p<0.05$). For log transformed counts see Figure 5.

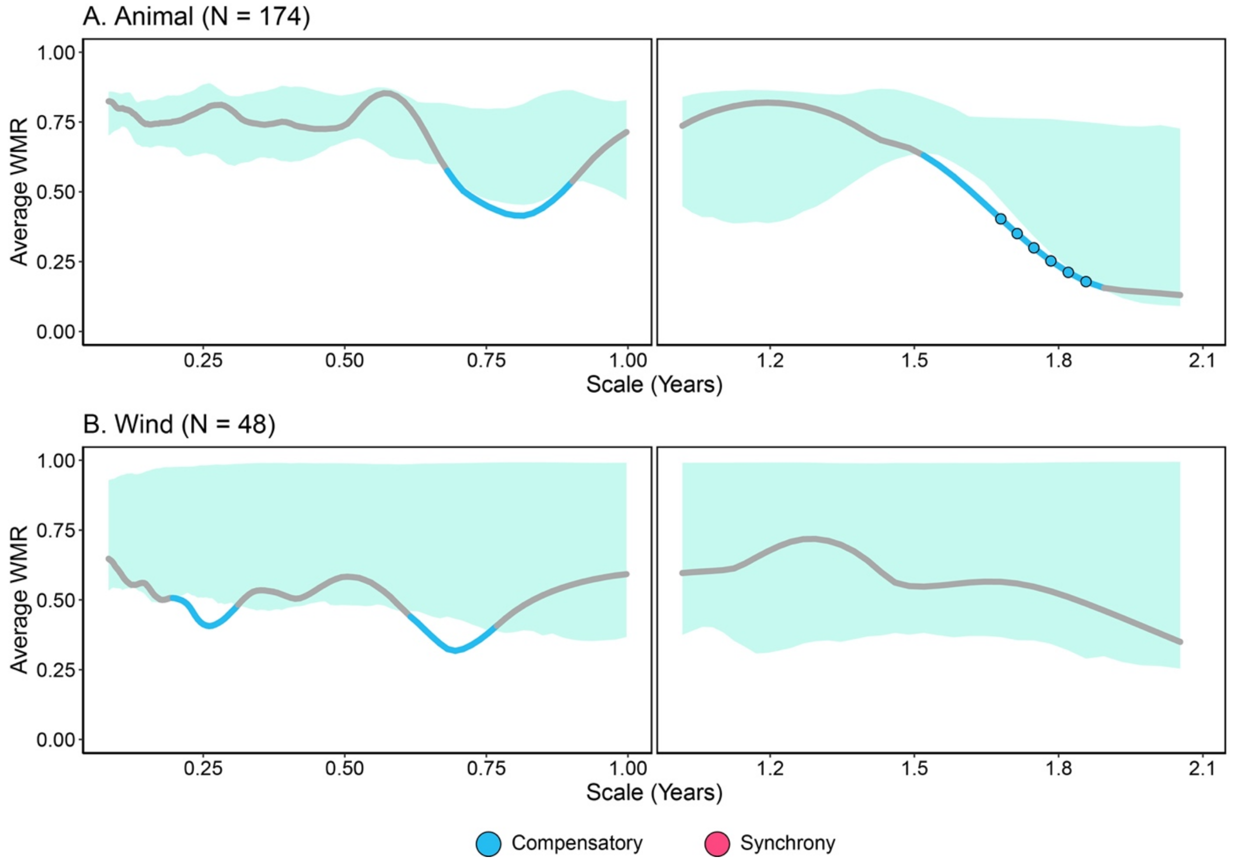

**Figure S11.** The averaged wavelet modulus ratio for tree species based on dispersal syndrome (thick lines), here showing raw untransformed counts for Cocha Cashu. The number in parentheses represents the number of species within the animal and wind-dispersed groups. The light blue ribbon represents the 2.5-97.5$^{th}$ percentiles of the null-distribution generated through bootstrapping. Any points that lie above the ribbon were considered nominally significant and synchronous while any points below the ribbon indicated nominally significant, compensatory dynamics ($p<0.05$). For log transformed counts see Figure S8.

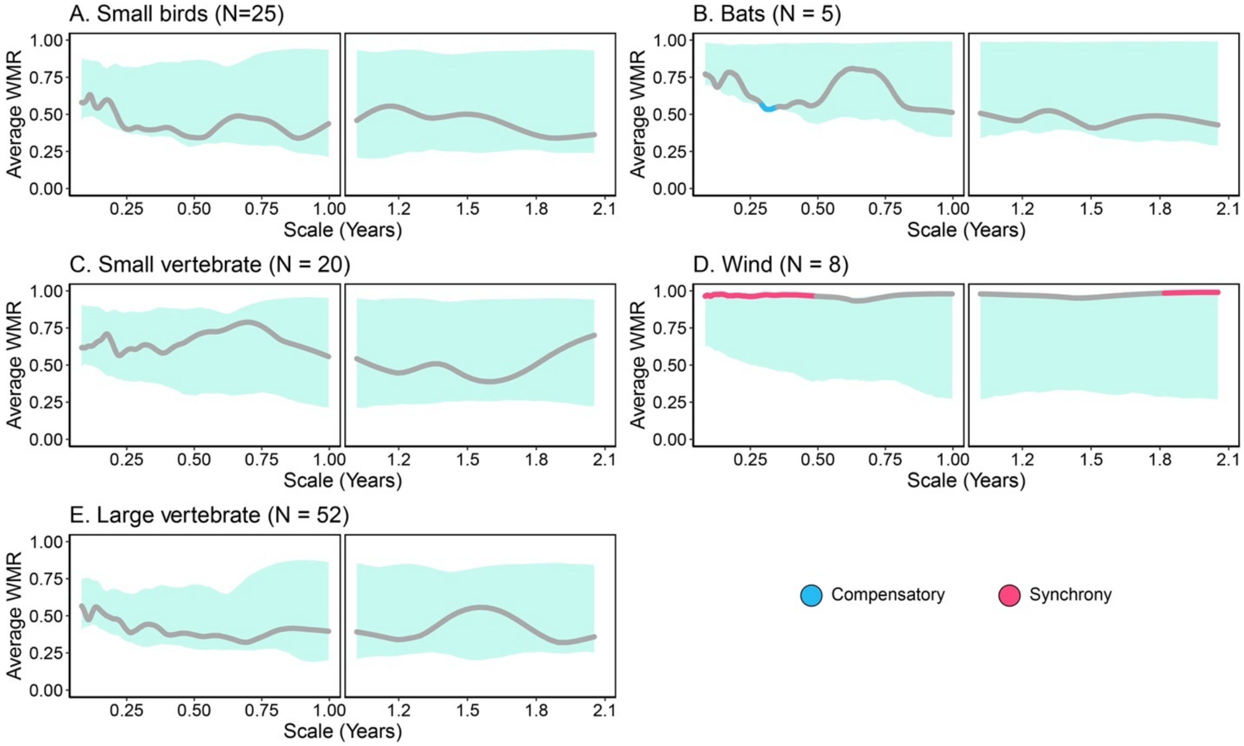

**Figure S12.** Wavelet power analysis (left panels) and average monthly wind (right panels) for the wind station data near Yasuní (1995-2012). Two wind parameters are shown: proportion of time calm (vs windy, top panels) and the average wind speed (bottom). Missing values were imputed using a weighted moving average with k = 2 in the imputeTS package in R.

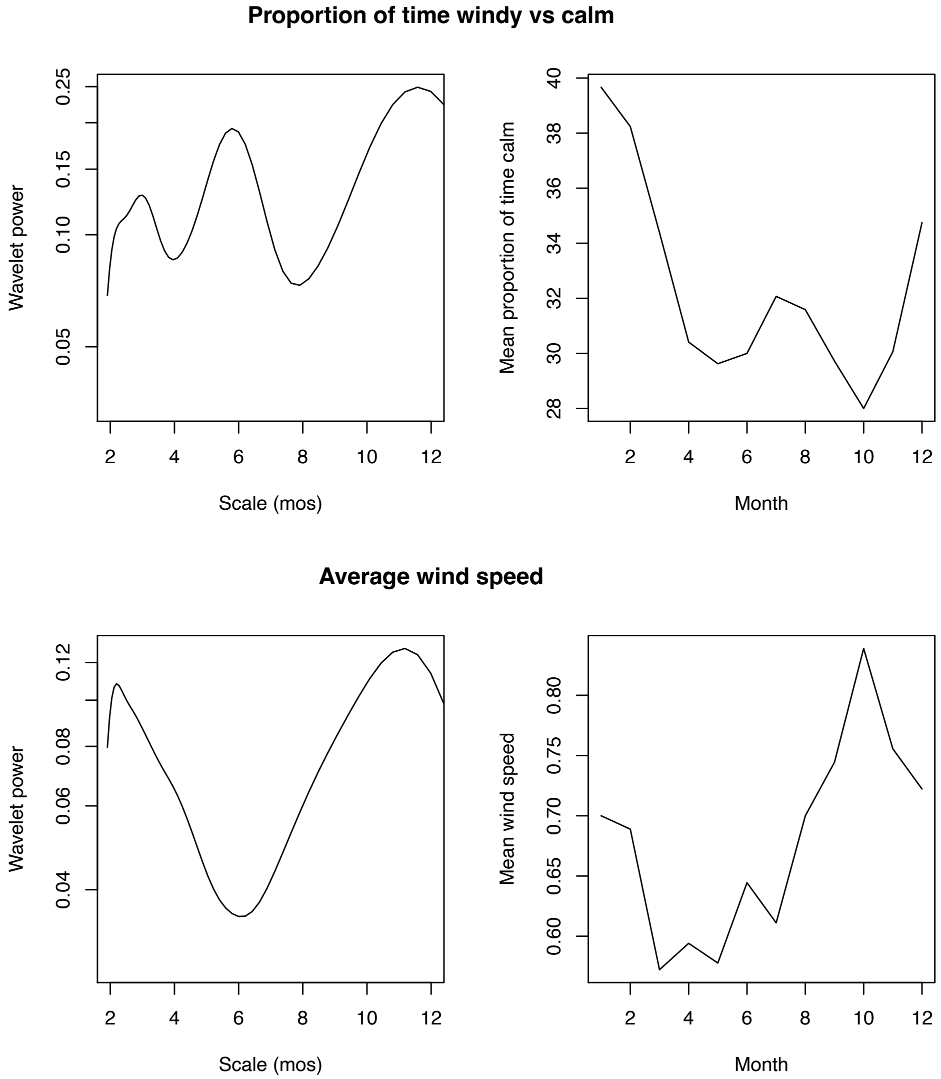

**Figure S13.** (A,C) Wavelet power across time scales for minimum temperatures, with higher power for a given scale indicating greater variability. (B,D) The Pearson correlation coefficient at varying scales (months) between the wavelet modulus ratio (WMR) of log transformed seed counts and wavelet-transformed climate. Positive correlations indicate that increases in the climate variable are associated with greater whole community synchrony, while negative correlations indicate the climate variable is associated with weaker whole community synchrony (or greater compensatory dynamics). The vertical lines represent the 2.5-97.5$^{th}$ percentiles of the null distribution (black circle shows mean of null) through phase-randomization permutation. Any points outside the distribution are considered nominally significant (colored in green), with those significant with FDR = 0.05 shown in green triangles.

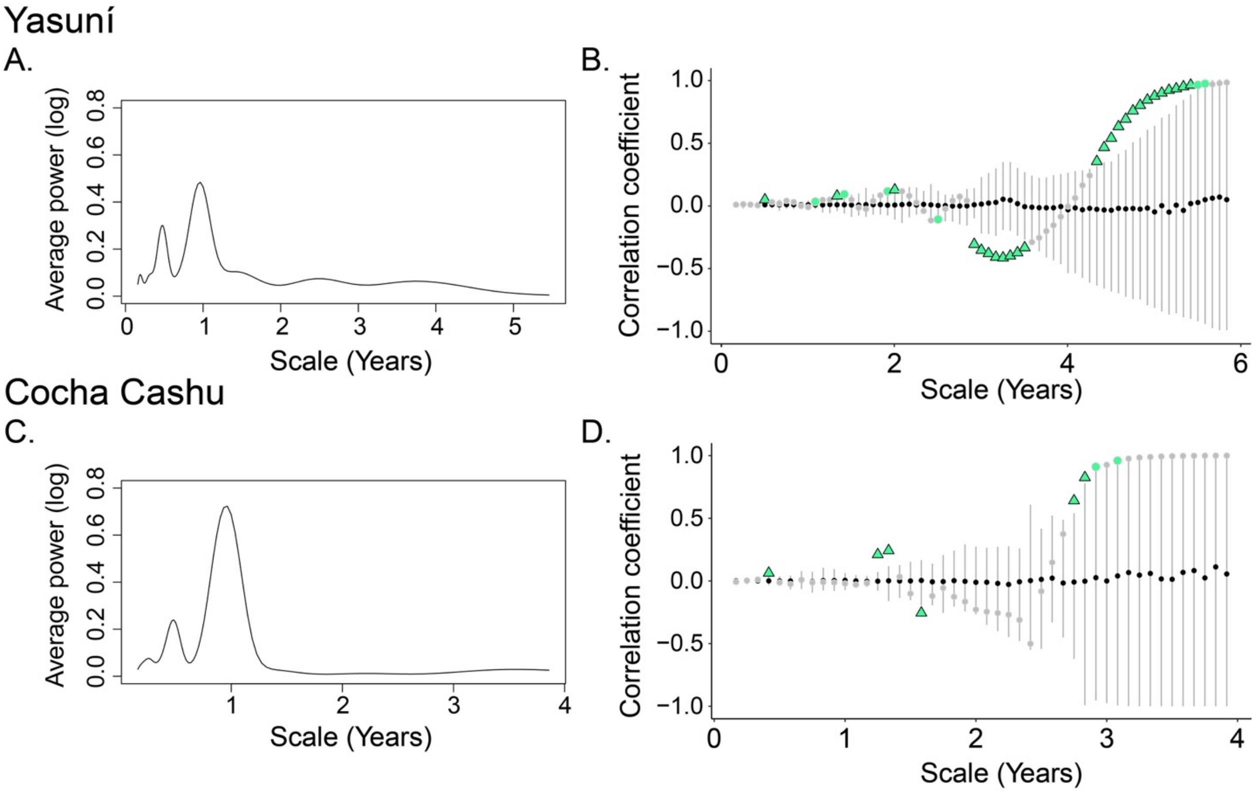

**Figure S14.** (A,C) Wavelet power across time scales for precipitation, with higher power for a given scale indicating greater variability. (B,D) The Pearson correlation coefficient at varying scales (months) between the wavelet modulus ratio (WMR) of log transformed seed counts and wavelet-transformed climate. Positive correlations indicate that increases in the climate variable are associated with greater whole community synchrony, while negative correlations indicate the climate variable is associated with weaker whole community synchrony (or greater compensatory dynamics). The vertical lines represent the 2.5-97.5$^{th}$ percentiles of the null distribution (black circle shows mean of null) through phase-randomization permutation. Any points outside the distribution are considered nominally significant (colored in green), with those significant with FDR = 0.05 shown in green triangles.

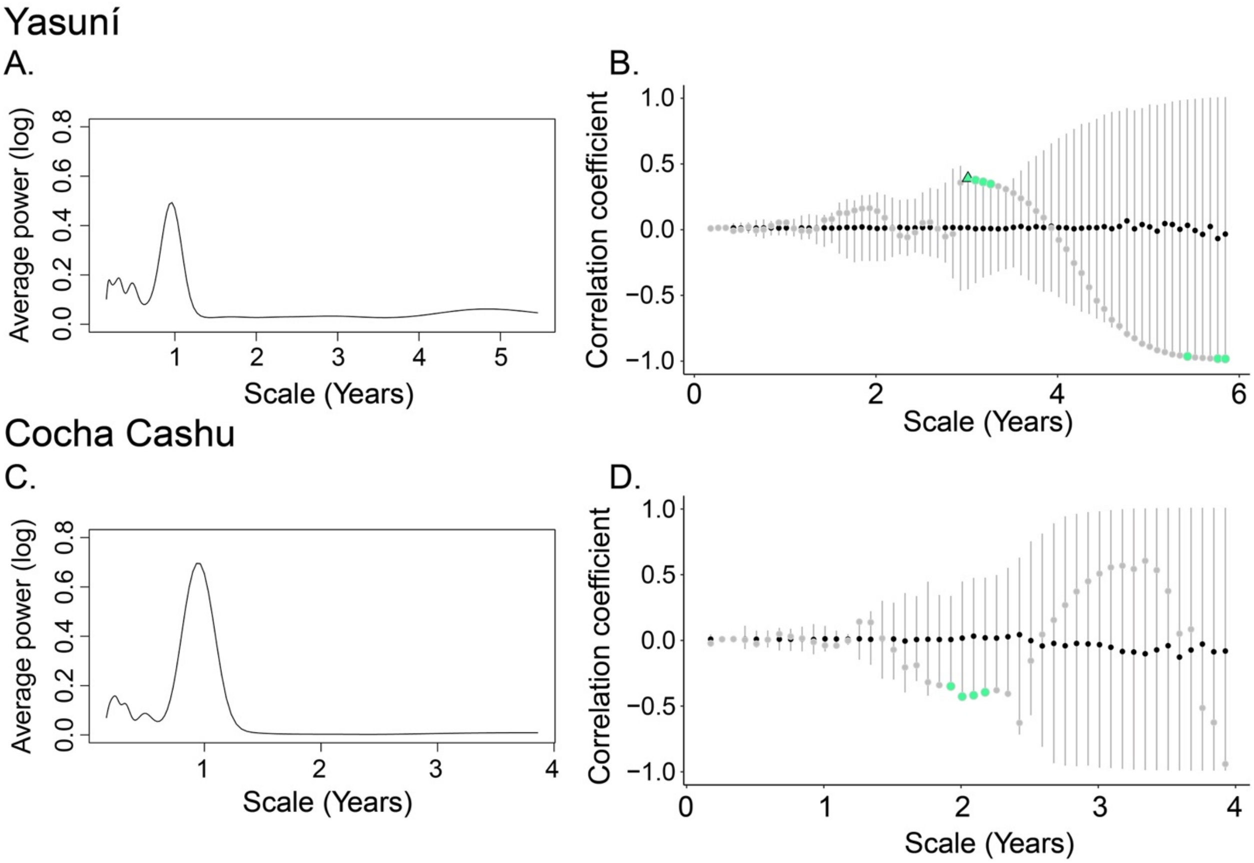

**Figure S15.** (A,C) Wavelet power across time scales for minimum temperatures, with higher power for a given scale indicating greater variability. (B,D) The Pearson correlation coefficient at varying scales (months) between the wavelet modulus ratio (WMR) of raw untransformed seed counts and wavelet-transformed climate. Positive correlations indicate that increases in the climate variable are associated with greater whole community synchrony, while negative correlations indicate the climate variable is associated with weaker whole community synchrony (or greater compensatory dynamics). The vertical lines represent the 2.5-97.5$^{th}$ percentiles of the null distribution (black circle shows mean of null) through phase-randomization permutation. Any points outside the distribution are considered nominally significant (colored in green), with those significant with FDR = 0.05 shown in green triangles.

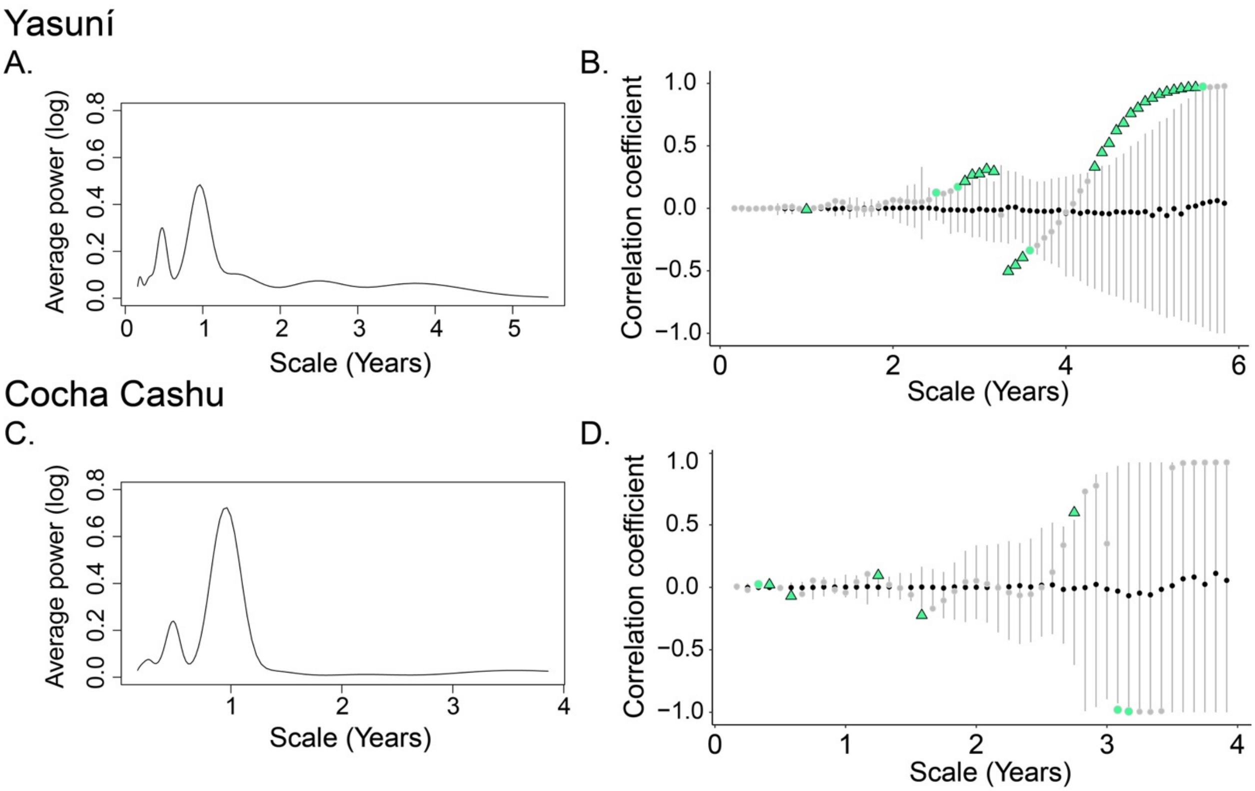

**Figure S15.** (A,C) Wavelet power across time scales for precipitation, with higher power for a given scale indicating greater variability. (B,D) The Pearson correlation coefficient at varying scales (months) between the wavelet modulus ratio (WMR) of raw untransformed seed counts and wavelet-transformed climate. Positive correlations indicate that increases in the climate variable are associated with greater whole community synchrony, while negative correlations indicate the climate variable is associated with weaker whole community synchrony (or greater compensatory dynamics). The vertical lines represent the 2.5-97.5$^{th}$ percentiles of the null distribution (black circle shows mean of null) through phase-randomization permutation. Any points outside the distribution are considered nominally significant (colored in green), with those significant with FDR = 0.05 shown in green triangles.

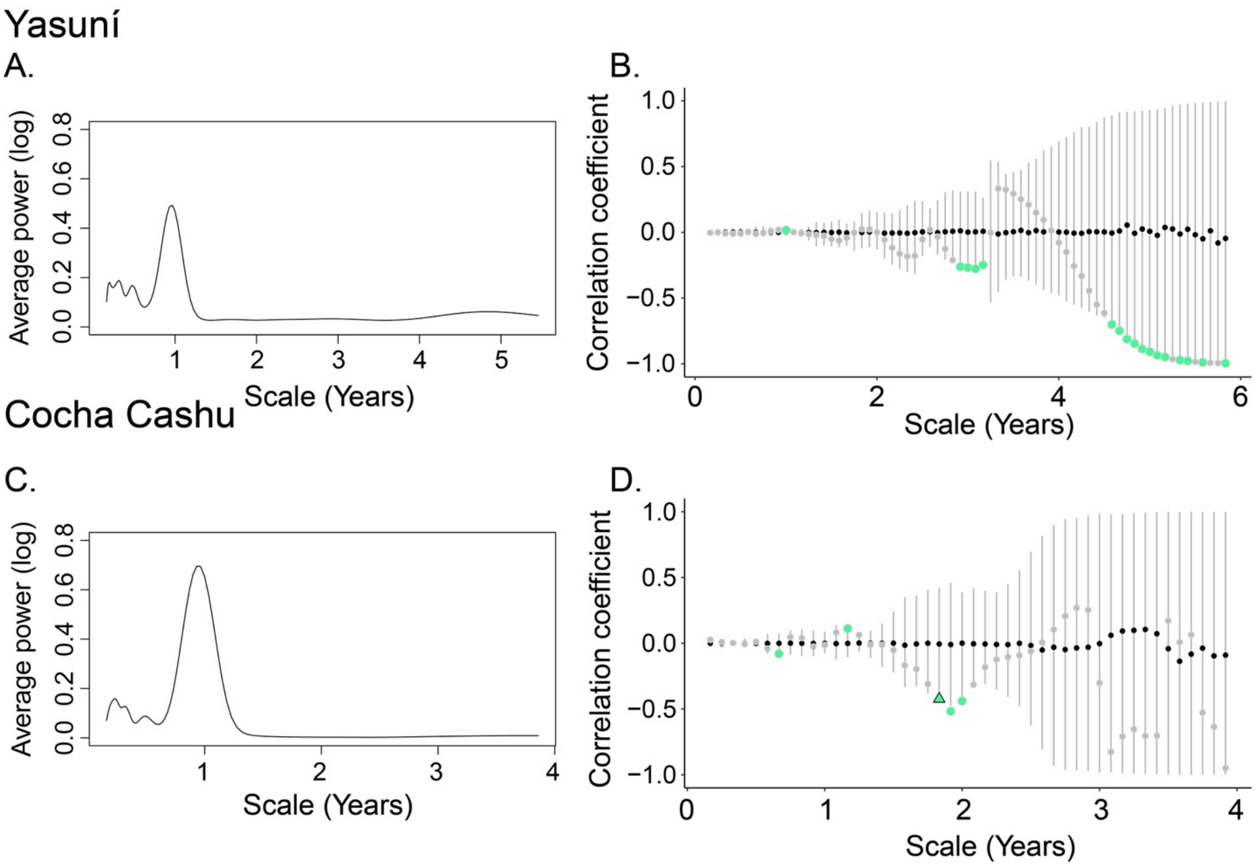

**Figure S16**. Yasuní monthly minimum temperatures averaged over 2000-2013, using ECMWF/ERA-Interim reanalysis data.

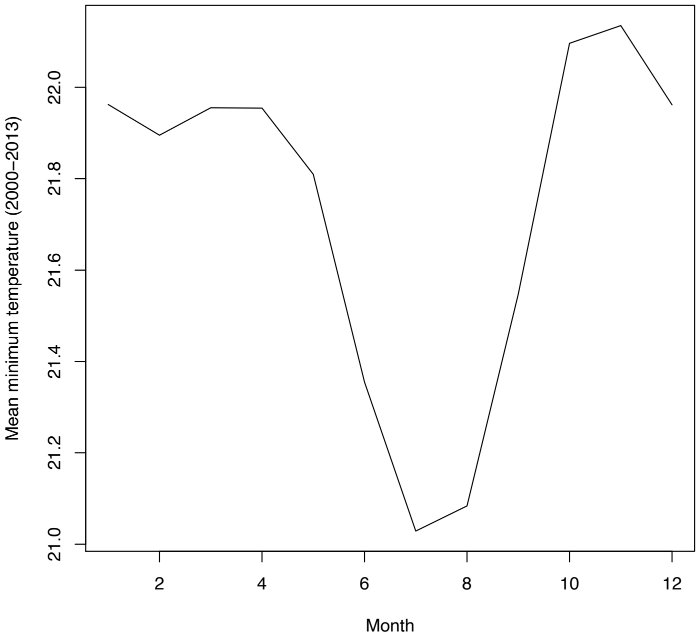

**Figure S17.** Whole community WMR on log transformed counts for Yasuní, excluding species with >95% dates with zero counts, resulting in only 330 species. The pattern is essentially the same as in Figure 2.

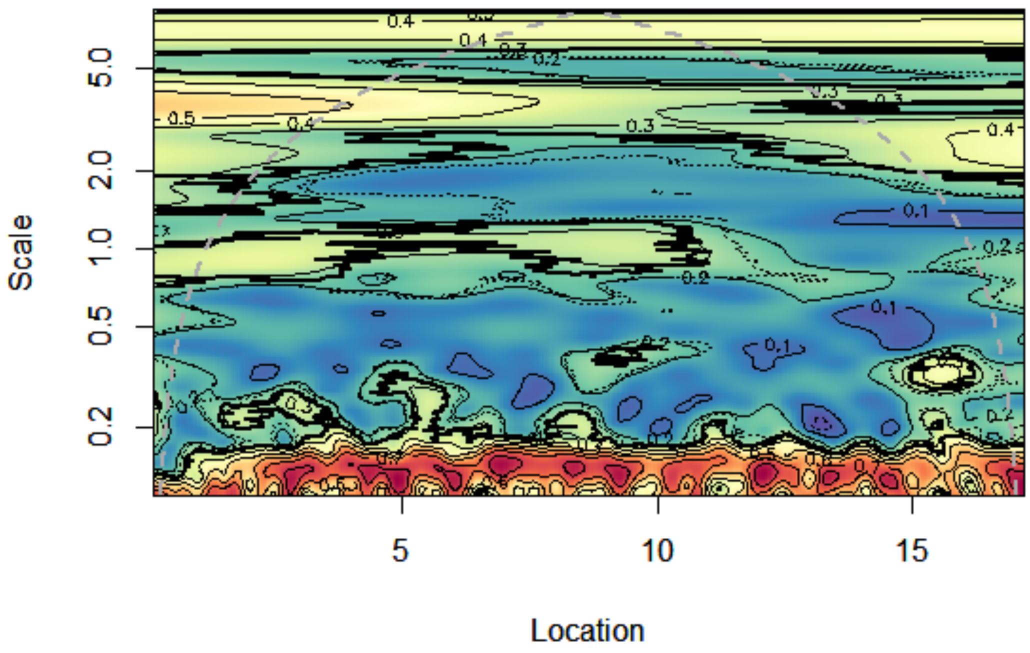

**Figure S18**. Whole community WMR on raw untransformed counts for Yasuní, excluding species with >95% dates with zero counts, resulting in only 330 species. The pattern is essentially the same as in Figure S2.

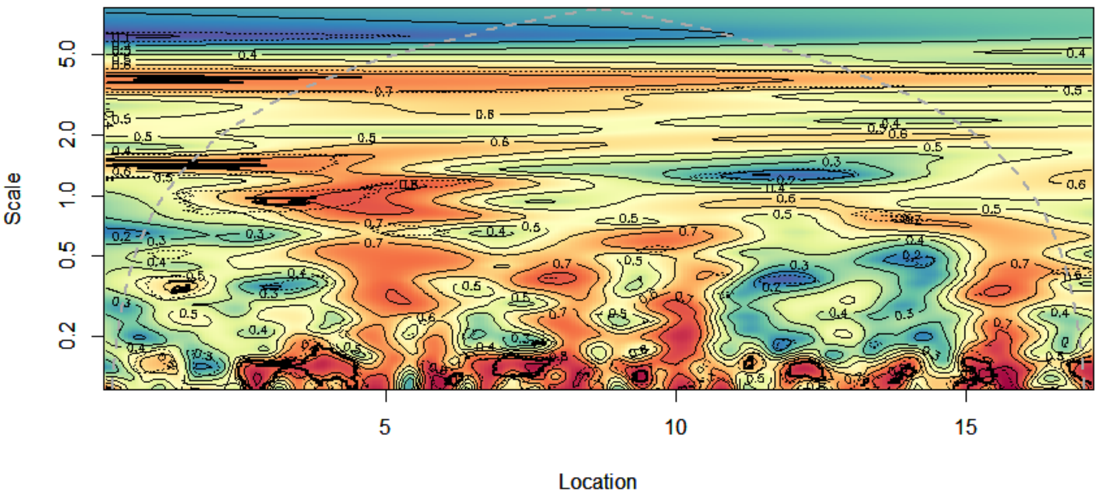

**Figure S19**. Whole community WMR on raw untransformed counts for Yasuní, excluding species with >75% dates with zero counts, resulting in only 66 species. The pattern is essentially the same as in Figure S2 and S18.

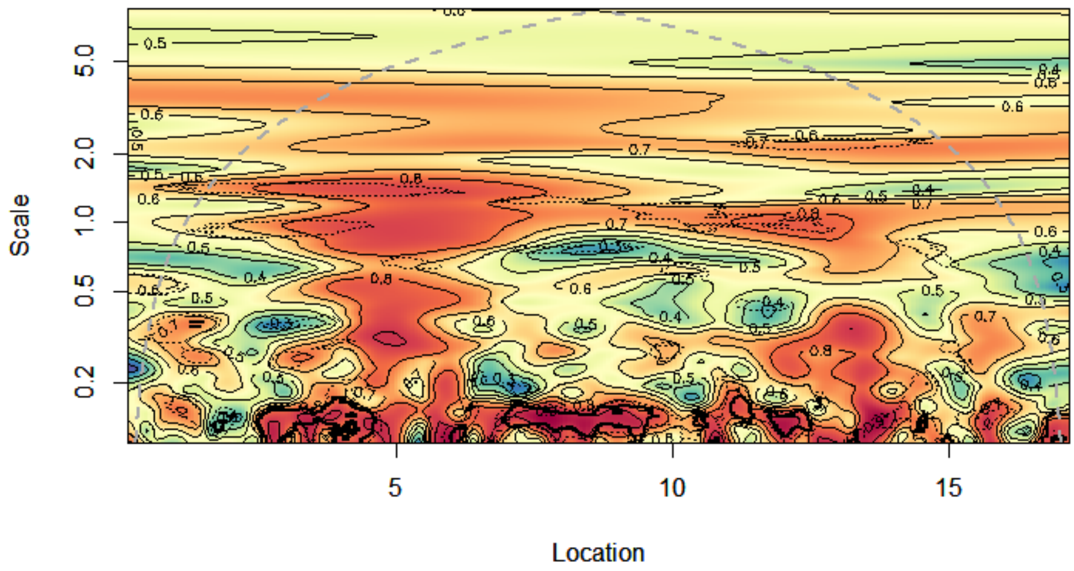

**Figure S20.** An example of a rare species at Yasuní, only observed on one interval in the middle of the time series, and the log($x$ + 1) transformation of its seed counts across scales. The transforms in the bottom three panels accurately capture the scale and location of the seed count variation (top panel). Compare to Figure S21 showing log($x$ + 0.1).

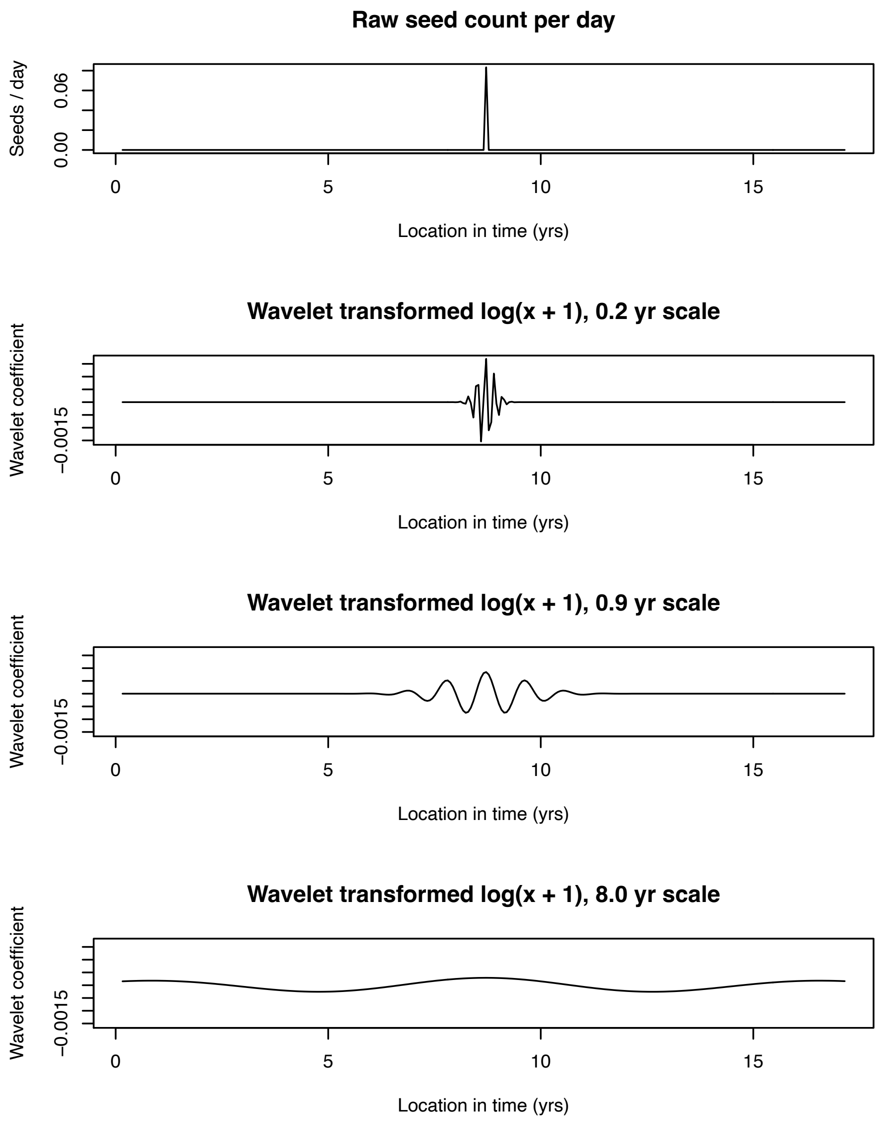

**Figure S21.** An example of a rare species at Yasuní, only observed on one interval in the middle of the time series, and the log($x$ + 0.1) transformation of its seed counts across scales. When counts are zero (as in nearly the whole time series), the log($x$ + 0.1) transformation gives non-zero values, and the wavelet basis function's variation approximately the only variation transmitted to the wavelet coefficient. This is problematic because the wavelet coefficients (bottom three panels) do not accurately capture the scale and location of the seed count variation (top panel). Compare to Figure S20 showing log($x$ + 1). The same problem occurs with log($x$ + 10).

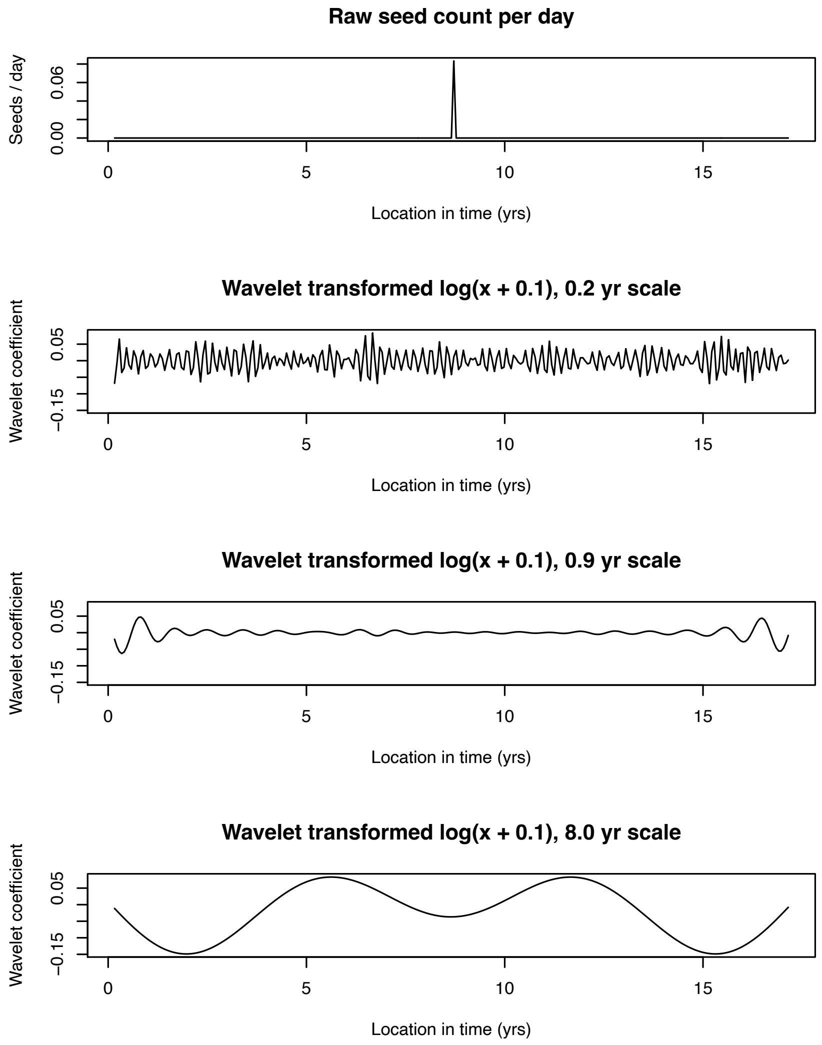

**Figure S22.** A further illustration of the problem of using different values (i.e. not 1) for *c* in the transformation of counts *x*, where we use log($x + c$) in our WMR calculations. Here Yasuní is shown. Significant synchrony is observed at all points in time and scales, due to most species' wavelet coefficients being dominated artifacts from the wavelet basis function.

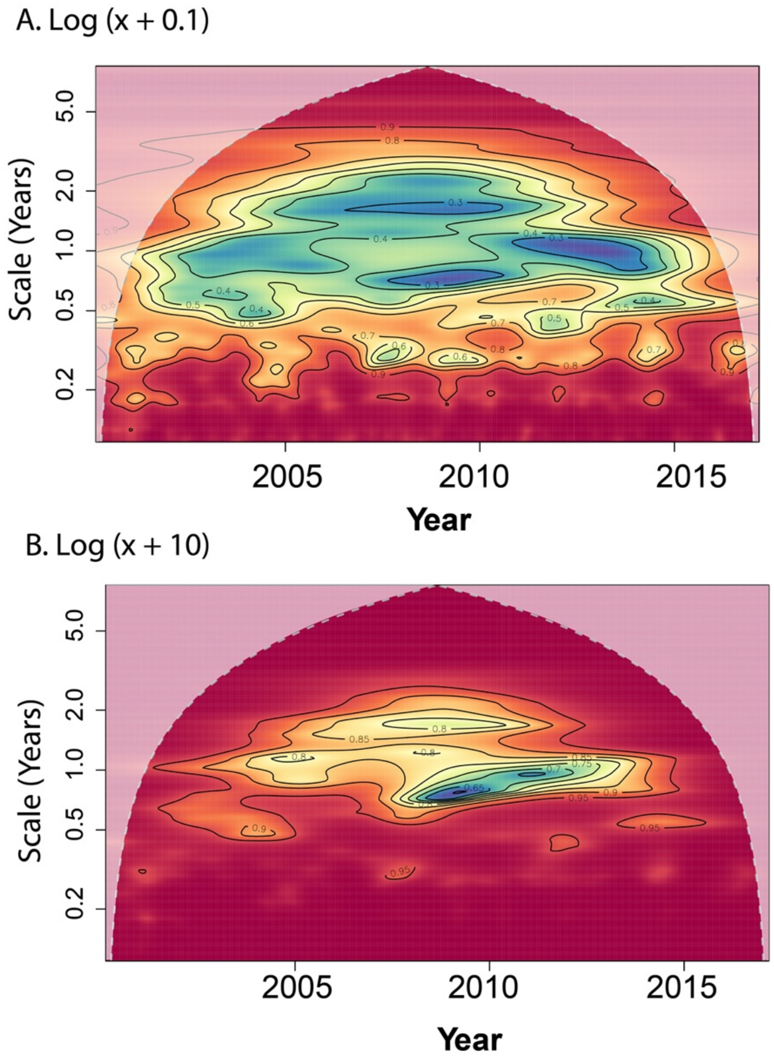

**Figure S23.** A further illustration of the problem of using different values (i.e. not 1) for *c* in the transformation of counts *x*, where we use log($x + c$) in our WMR calculations. Here Cocha Cashu is shown. Significant synchrony is observed at nearly all points in time and scales, due to most species' wavelet coefficients being dominated artifacts from the wavelet basis function.

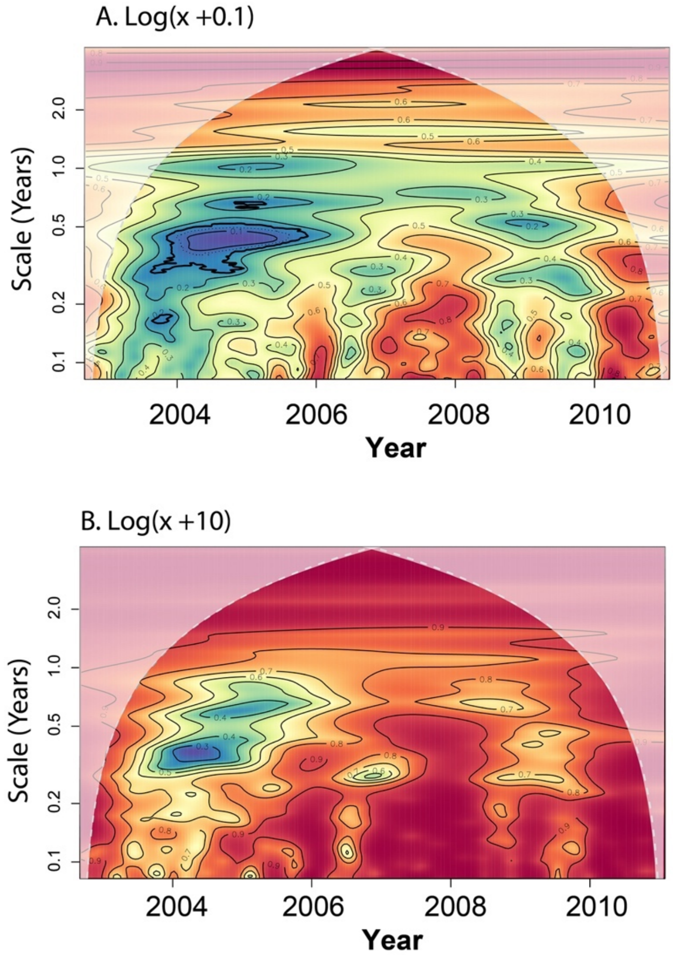
